# Single-Cell and Spatial Transcriptomic Analyses Reveals the Dynamic Transcript Profiles of Myocardial Lymphangiogenesis post Myocardial Infarction

**DOI:** 10.1101/2024.06.10.598235

**Authors:** Jiaqi He, Dali Zhang, Haixu Song, Ziqi Liu, Dan Liu, Xiaolin Zhang, Xiaojie Zhao, Yan Zhang, Jing Liu, Jiaxin Xu, Chenghui Yan, Yaling Han

## Abstract

Cardiac lymphatics play an important role in myocardial edema and inflammation. This study integrated single-cell sequencing data and spatial transcriptome data from mouse heart tissue at different time points post-myocardial infarction (MI), and identified four transcriptionally distinct subtypes of lymphatic endothelial cells (LECs) and localized them in space. Interestingly, LECs subgroups was found to be localized in different zones of infarcted heart related to different functions. Additionally, LEC capillary III(LEC ca III) may be involved in the direct regulation of myocardial injuries in infarcted zone from the perspective of metabolic stress, while LEC ca II may be related to the rapid immune inflammatory responses of the border zone in the early stage of MI. LEC ca I, as well as LEC collection mainly participate in the regulation of myocardial tissue edema resolution in the middle and late stages post-MI. Cell trajectory and Cell-Chat analyses further identified that LECs may regulate myocardial edema through Aqp1, and might affect the infiltration of macrophages through the Galectin9-CD44 pathway. Collectively, our study revealed the dynamic transcriptional heterogeneity distribution of LECs in different regions of the infarcted heart, in detail; these LECs formed different functional subgroups, that might exhibit different bioeffects in myocardial tissue post-MI.

## 1 Introduction

The lymphatic vasculature, which accompanies the blood vasculature in most organs, is indispensable for maintaining of tissue fluid homeostasis, immune surveillance and lipid uptake and transport(Alitalo, Tammela, & Petrova, 2005; Huang, Elvington, & Randolph, 2015; Petrova & Koh, 2020). In contrast to circulatory blood vessels, lymphatic vessels are structurally blind-ended segments, that are designed to absorb interstitial fluid and immune cells. These lymphatic capillaries converge onto deeper lymphatic structures called collecting vessels, which are equipped with valves and a muscular wall that promote unidirectional transport and drain extravascular fluid, macromolecules and immune cells(Alitalo, 2011; Oliver, Kipnis, Randolph, & Harvey, 2020).

The cardiac lymphatic vasculature was first described in the 17^th^ century(Tammela & Alitalo, 2010). To date, studies have demonstrated the role of the lymphatic system only in embryonic heart development and cardiac health in adults(Brakenhielm & Alitalo, 2019; Heron, Ratajska, & Brakenhielm, 2022; Klotz et al., 2015). Cardiac lymphatics can take up inter-tissue fluid and transport immune cells through highly branched, blunt, open-ended lymphatic capillaries, to the lymph nodes near the thoracic ducts and, finally, into the superior vena cava(Huang, Lavine, & Randolph, 2017).

Myocardial infarction (MI) is one of the most important cardiovascular diseases. Myocardial edema and inflammation are the two dominant pathological processes in the early stages of MI(Anzai, Ko, & Fukuda, 2022; Bulluck, Dharmakumar, Arai, Berry, & Hausenloy, 2018; Chen, Zhao, & Yan, 2022; Nahrendorf, Pittet, & Swirski, 2010). Many preclinical studies have revealed severe remodeling of lymphatic system during MI in animal models(Ishikawa et al., 2007; Shimizu et al., 2018; Vuorio et al., 2018). The stimulation of cardiac lymphangiogenesis by adrenomedullin or vascular endothelial growth factor C can ameliorate myocardial edema following MI(Henri et al., 2016; Trincot et al., 2019). However, owing to the extremely low number of lymphatic vessels in the myocardial tissue, there is still a lack of comprehensive and in-depth understanding of the pathological mechanisms underlying myocardial injury and prognostic progression after MI.

With the rapid development of single-cell transcriptome sequencing and spatial single-cell transcriptome sequencing technologies, researchers can explore cell subpopulations, heterogeneity, function, metabolism, and cell-cell interactions at the single-cell level(Grün & van Oudenaarden, 2015; Vandereyken, Sifrim, Thienpont, & Voet, 2023). In order to further elucidate the effects and regulatory mechanisms of the lymphatic vessels in the repair process of myocardial injury following MI, this study integrated single-cell sequencing and spatial transcriptome data from mouse heart tissue at different time points after MI from publicly available data (E-MTAB-7895, GSE214611) in the ArrayExpress and gene expression omnibus (GEO) databases. By reanalyzing these data, this study revealed dynamic transcriptional heterogeneity and spatial localization differences in regenerated lymphatic endothelial cells (LECs) in heart tissue after MI. Different subpopulations of LECs appear at different times and locations during the process of myocardial post-MI, and perform different biological functions. Furthermore, we validated two key regulatory molecules that play important regulatory roles in myocardial edema and lymphatic immune regulation after MI.

## 2 Results

### 2.1 Dynamic Changes of Cardiac Lymphatics in Cardiac Tissue post-MI

To explore the physiological and pathological functions of cardiac lymphatic vessels after myocardial infarction (MI), we established an MI mouse model by ligating the anterior descending branch of the coronary artery (n=32) (Figure1-figure supplement 1A). Heart function was evaluated by micro-ultrasound, and mouse hearts were harvested at 0, 3, 7, 14, and 28 days after MI (n=5-7/ per time-point) (Figure1-figure Supplement 1B). Immunofluorescence (IF) staining was performed using antibodies against Prospero homeobox 1 (Prox1, nuclear protein), the lymphatic vessel endothelial hyaluronan recetor-1 (Lyve1, membrane protein) and podoplanin(Pdpn, membrane protein), all of which specifically labeled the LECs (Figure1A, Figure1-Figure Suppl-1C-E). Since the membrane protein Lyve1 and Pdpn can present lymphatic vessel morphology more clearly than Prox1, Lyve1 and Pdpn were used for lymphatic vessel co-staining at 0, 3, 7, 14, and 28 days after MI (Figure 1A, Figure1-Figure Suppl-2A). There were significant differences in the morphology and distribution of the newly formed lymphatic vessels in different regions of the heart tissue post-MI (Figure 1B). Three days post-MI, lymphatic vessels in the infarcted zone (IZ) and border zone (BZ) were significantly lost. However, on the 7th day, a large number of newly formed and dilated continuous lymphatic vessels appeared in the BZ. In the IZ, only a small number of lymphatic capillaries with discontinuous walls were observed. On the 14th and 28th days following the infarction, the number of lymphatic capillaries in the IZ significantly increased. Continuous large lymphatic collecting vessels were observed in the epicardium of the IZ, and the number of large lymphatic collecting vessels in the BZ did not decrease (Figures 1A and 1B, Figure1-figure Supplement 2B). In addition, there were no significant changes in the distribution and morphology of cardiac lymphatic vessels in the unaffected normal myocardial area (remote zone, RZ) at any time (Figure1-figure Supplement 2A). These results suggest that lymphatic regeneration may be involved in the regulation of myocardial damage following MI.

**Figure 1.**
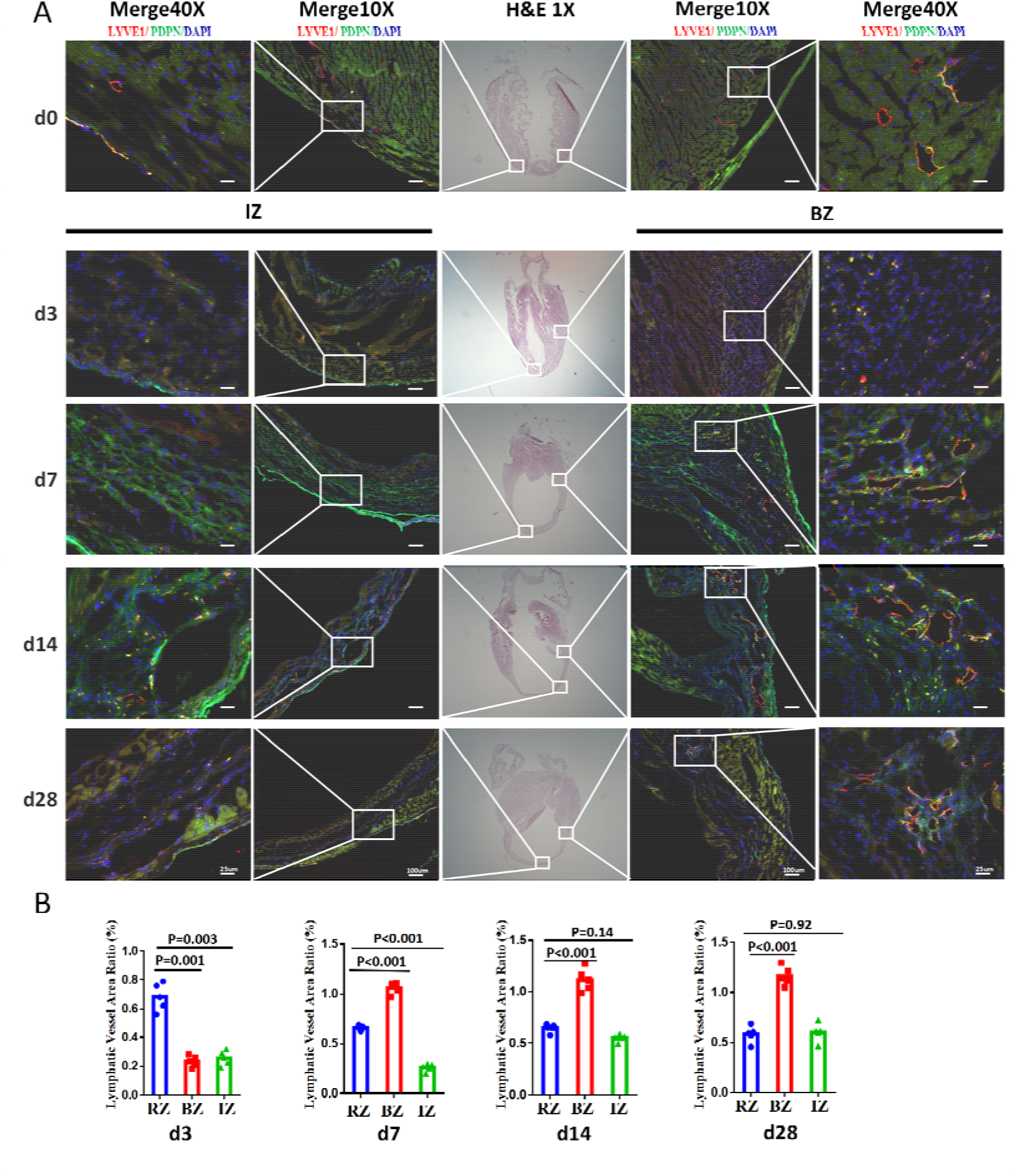
Dynamic Changes of Cardiac Lymphatics in Cardiac Tissue after MI. **(A)** Hematoxylin-eosin (H&E) staining and immunofluorescence (IF) co-staining with LYVE1 and PDPN of tissue sections in sham and MI mice model at d3, d7, d14, and d28 post-MI (n=5∼7/group). LYVE1: lymphatic vessel endothelial hyaluronan receptor 1; PDPN: Podoplanin; DAPI: 4’6-diamidino-2-phenylindole; IZ: Infarct Zone; BZ: Border Zone; the scale bars are in 10**×**-100μm and 40**×**-25μm;(**B**)Semi-quantitative fluorescence determination of Lymphatic vessel area ratio under 100-fold magnification by ImageJ(n=5).

### 2.2 Identification of Four Transcriptionally Distinct Subtypes of Regenerated Lymphatic Endothelial Cells in Myocardial Tissue post-MI via Single-Cell Sequence

To further clarify why the regenerated cardiac LECs mapped to a particular distribution during cardiac repair post-MI, a set of published single-cell sequencing data (E-MTAB-7895) was selected from the ArrayExpress database of mouse MI models at different time points (days 0, 1, 3, 7, 14 and 28). Using specific marker proteins as cell-clustering markers, the cluster dimensionality reduction analysis of the 44, 839 single cells (excluding myocardial cells) in this experimental dataset was recalculated. First, we classified these cells into five categories based on specific genes: endothelial cells (Cdh5, Pecam1, and Vwf), fibroblasts (Dcn, Col1a1, and Gsn), immune cells (Ptprc, and C1qa), myocardial fibroblasts (Cthrc1, and postn), smooth muscle cells and pericytes (Myh11, Tagln, Acta2, Rgs5, Abcc9, and Kcnj8) (Figures. 2A and 2B). Subsequently, the endothelial cells (Cdh5, Pecam1, and Vwf) were further clustered into five sub-clusters, including angiogenic endothelial cells (Col4a2, Apln, Sparcl1, and Aplnr), arterial endothelial cells (Fbln5, Hey1, Sema3g, and Efnb2), capillary endothelial cells (Kdr, Rgcc, Aqp1, and Endou), venous endothelial cells (Mgp, Cfh, Bgn, Vwf, and Plvap), and LECs (Lyve1, Cldn5, and Prox1) (Figures. 2C and 2D). To further characterize LECs, we examined the expression of Foxp2, Ackr4, Cavolin-1(Cav1), Nt5e, Pdpn and Lyve1(Takeda et al., 2019). Consistent with this report, the collecting LEC marker Cav1 was expressed in the clusters LEC collecting (LEC co), LEC capillary I (LEC ca I), and LEC capillary III (LEC ca III) clusters, whereas Foxp2 and Ackr4 were expressed almost exclusively in LEC co clusters. Capillary markers Pdpn and Lyve1 were expressed in four subsets, but showed the highest expression in LEC ca III, and the lowest in LEC co (Figures. 2E and 2F). Together, these results indicate significant transcriptional heterogeneity in the regenerated cardiac LECs post-MI.

**Figure 2.**
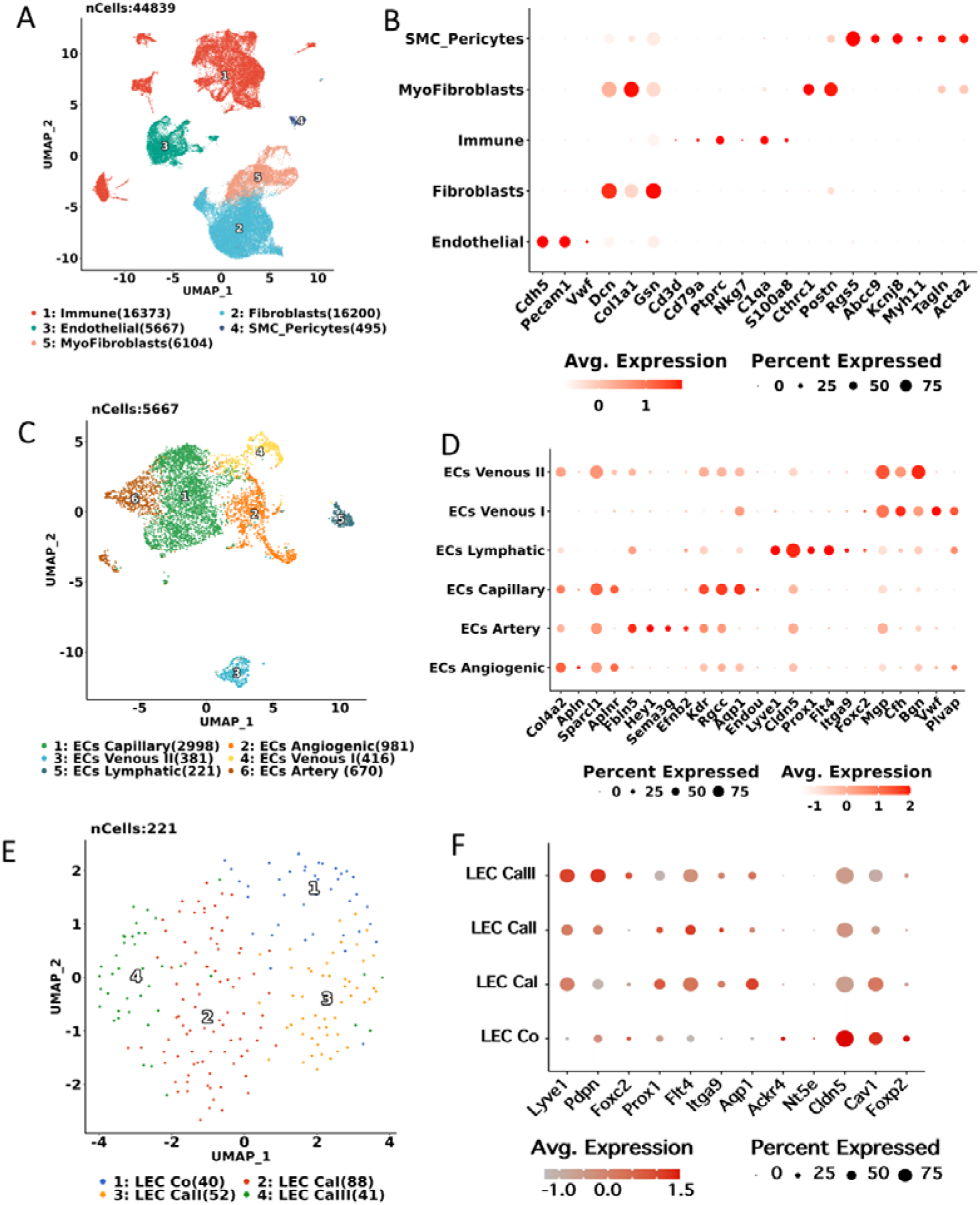
Identifying infiltrated cell types in tissues. **(A)**. A Uniform Manifold Approximation and Projection (UMAP) plot of 44,860 showing five identified non-myocardium cell types in infarcted heart, cell types are coded with different colors; **(C)**. A UMAP plot of single cells profiled in the presenting work colored by endothelial cell types; **(E)**. A UMAP plot of single cells profiled in the presenting work colored by LEC cell types; **(B, D, and F)**. Dot plot showing the average expression of selected cell marker genes. The dots’ size and color indicate the percentage of marker gene expression and the average expression level (Avg. Expression).

### 2.3 Relationship Between the Spatial Localization and Function of LEC Subpopulations post-MI

To elucidate the biological functions of LECs subgroups in myocardial tissue post-MI, we first observed dynamic changes in the number and proportion of LECs subsets after MI. Consistent with the results of IF staining, the number of LECs was significantly reduced in the early phases of MI (D1 and D3). From D7, the number of cells gradually increased and returned to normal levels within 14 days. The four LECs subtypes differ in their proliferation potential, with CaI and CaIII appearing to proliferate faster than Co and CaII. Normal myocardial tissues are mainly composed of LEC ca I, LEC ca II, and LEC co, with a small amount of LEC ca III. In the early stages of myocardial infarction (D1 and D3), the number of LECs decreased sharply. The number of LECs gradually increasing from day 7 and returning to normal levels by day 14 after MI. Moreover, from day 14 onwards, the number and proportion of Ca I type LECs significantly increased. Twenty-eight days post-MI, the number and proportion of the three capillary LECs had returned to normal, except for a small proportion of LEC co in the group (Figures. 3A and 3B).

**Figure 3.**
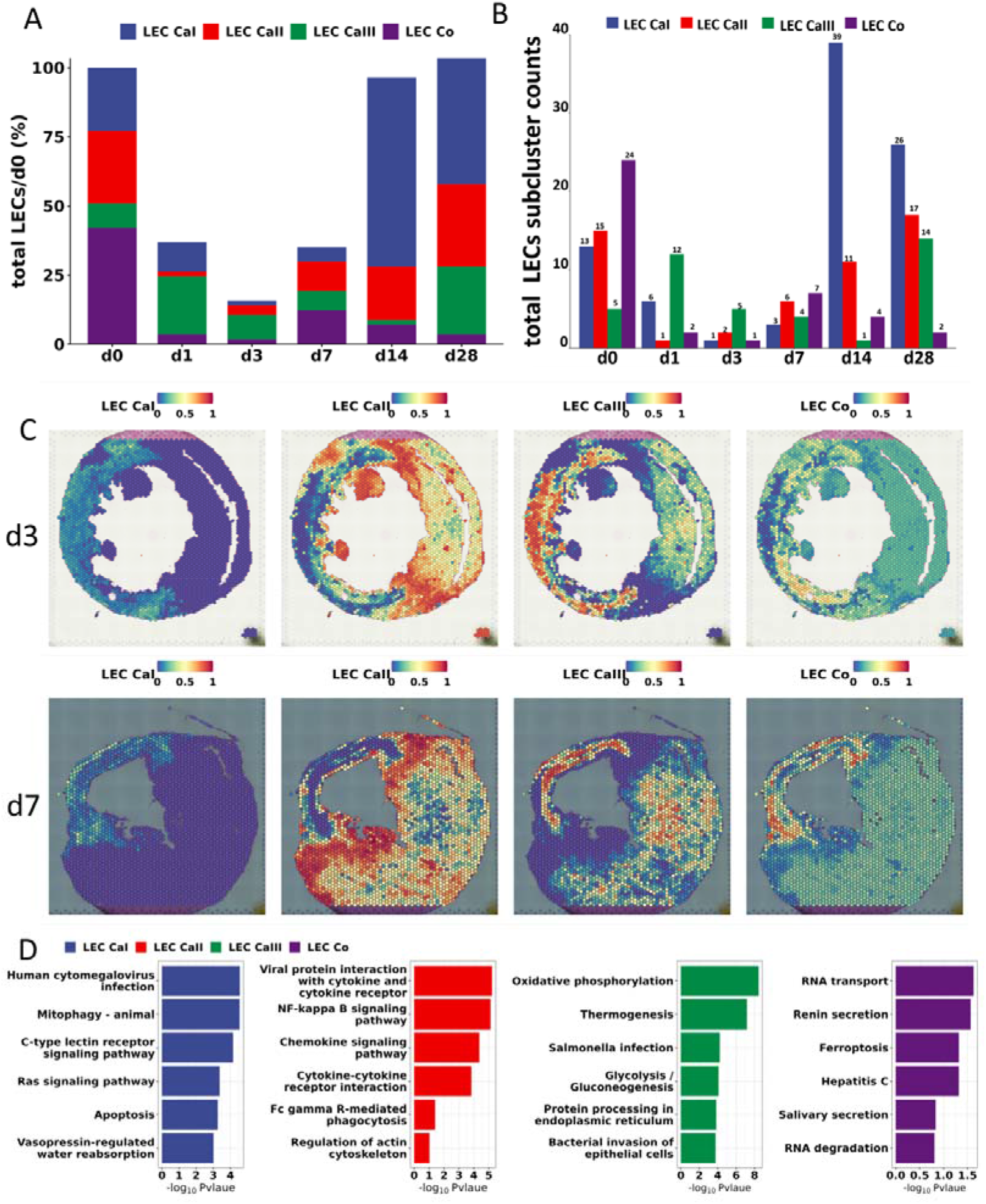
The Relationship Between the Spatial Localization and Function of LEC Subpopulations post MI. **(A)** Bar plots showing total numbers and subtypes of LECs at different time points post-MI. The y-axis represents the percentage of the total number of LECs at d1, d3, d7, d14, and d28 post-MI, relative to the number of LECs at d0, which is used as the reference value set at 100%. **(B)** Bar plots showing total cell counts of each subcluster at different time points. **(C)** The proportion of LEC cell types in each spot determined by spatial transcriptome in d3 and d7 post-MI; **(D)** Enriched KEGG pathways of upregulated genes in different LEC cell types, color indicate different cell types.

To validate whether these LEC subpopulations are involved in the repair of myocardial injury following MI, we retrieved published single-cell nuclear transcriptomic and spatial transcriptomic data (GSE214611) of mouse MI models from the GEO database. In early MI (D3), LEC ca I, ca III and LEC co were mainly distributed in the IZ and BZ, with LEC ca III being significantly higher in the IZ. In contrast, LEC ca II was mainly distributed in the BZ and RZ, and was significantly reduced in the IZ. As the morphology of the infarcted area gradually became clear, on D7, LEC ca I continued to increase in the IZ region, LEC ca III was still mainly distributed in the IZ, LEC co migrated from the BZ to the IZ, and LEC ca II showed no significant spatial localization or expression differences, but were still highly expressed in the BZ and RZ (Figure 3C, Figure3-figure supplement 1).

KEGG functional enrichment analysis was conducted on the genes of the four LEC subtypes. LEC ca III, the LEC subgroup with the highest proportion expressed in the IZ during the early stage of MI, was found to primarily participate in energy metabolism such as oxidative phosphorylation, thermogenesis, glycolysis/gluconeogenesis, antigen presentation, cell morphology and migration related functions. Although ca I and ca III have similar spatial localization after MI, their expression levels were lower than that of LEC ca III on D3, but significantly increased in the IZ on D7. LEC ca I was found primarily participate in the activation of infection, apoptosis, mitophagy, and water absorption related regulation of vasopressin. LEC co showed partial similarity in biological functions with ca I, and were mainly involved in fluid regulation and flow, such as renin, saliva secretion, and inflammation. LECs are inconsistently is involved in the regulation of RNA transport, degradation, and ferroptosis. LEC ca II, which significantly reduced in IZ, while sustaining high expression in the BZ and RZ, is mainly related to NFκB is associated with the regulation of chemokine signaling pathways and cytokine interactions between ligands (Figure 3E).

In summary, based on the pathological changes in myocardial tissue, as well as the regeneration ratio, spatial localization, and functional differences of LEC subgroups after MI, we deduced that LEC ca III may be involved in the direct regulation of myocardial injury in the IZ from the perspective of metabolic stress, whereas LEC ca II may be related to the rapid immune inflammatory response in the BZ in the very early stages of MI. LEC ca I and LEC co primarily participate in the regulation of myocardial tissue edema resolution in the middle and late stages post-MI.

### 2.4 Single-Cell Trajectories of LECs Subsets post-MI

How do LEC subgroups regenerate and evolve after MI? To answer this question, we used cell trajectory analysis to characterize LECs development and determine lineage differentiation. From the pseudo-time plot and phenotype-based trajectory plot, we learn that, in the early stage after MI, the main LEC subgroup is still LEC ca II and III, which is consistent with the spatial transcriptome localization results. LEC ca II was significantly upregulated in the BZ and RZ of the infarcted area, and Ca III was significantly upregulated in the IZ. During the mid-to-late stages, we found that LEC ca II gradually differentiated into two orientations. One group was mainly composed of ca I, which may regulate immune and metabolic outcomes, while the other group jointly involved ca I, ca III, and co, which may be mainly involved in cell death, inflammation and edema regression (Figures. 4A-C). These results further suggest that ca II and III mainly participate in the early stages of myocardial infarction, whereas ca I and co are involved in the regulation of injury repair in the middle and late stages after MI.

**Figure 4.**
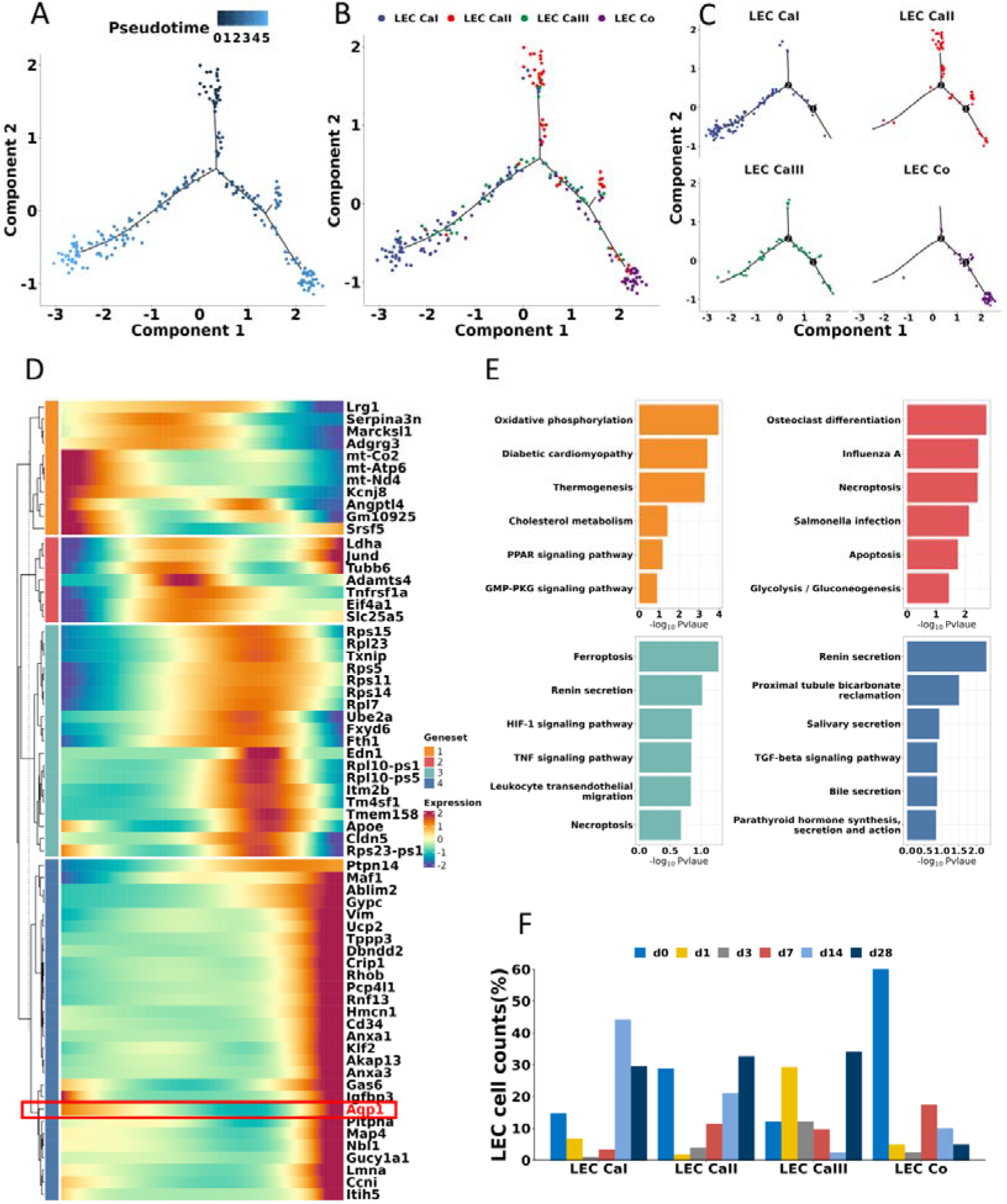
Single-Cell Trajectories of LECs Subsets post MI. **(A, B, and C)**. Pseudo-time analysis by Monocle2 shows the potential evolutionary trajectory of LECs, colored by cell types; **(D)**. Heatmap shows the set of genes with altered expression levels along the pseudo-time; **(E)** Enriched KEGG pathways in four different gene sets, color indicate different cell types; **(F)** Bar plots show the proportion of LECs subgroups at different time points.

What are the main regulatory molecules in these LEC subgroups during pseudo-time post-MI? Subsequently, we screened and performed KEGG functional enrichment analysis on temporally expressed genes during MI and obtained four gene sets (Figures. 4D and 4E). Interestingly, these genes corresponded well with the ratios of different LEC subgroups during MI (Figure 4F). Gene set 1, similar to LEC ca III, which is highly expressed at the very early stages of MI and then gradually decreases, is mainly related to oxidative phosphorylation, diabetic cardiomyopathy, cholesterol metabolism, thermogenesis, and the peroxisome proliferator activated receptor (PPAR), cyclic guanosine 3’, 5’-monophosphate (cGMP) - Protein Kinase G (PKG) signaling pathway. The expression of gene sets 2 and 3 was lower during MI, significantly increased in the early and middle stages, and gradually decreased thereafter. These genes are primarily involved in the regulation of programmed cell death, apoptosis, amino acid metabolism, glycolysis, and gluconeogenesis. Gene set 4 showed lower expression in the early stage of myocardial infarction, but gradually increased in the middle and showed the highest expression in the late stages post-MI. These genes are mainly involved in cardiac regeneration, renin secretion, bicarbonate reabsorption, salivary and bile secretion, as well as parathyroid and growth hormone synthesis and secretion. These results imply that LECs are mainly involved in the stress regulation of cell metabolism, inflammation, and death in the very early stages of MI, while regeneration and repair functions are mainly involved in the late stages.

### 2.5 Assessment of Cell-Cell Communication between LECs and immune cells

To explore how LECs and immune cells interact to regulate metabolic inflammation in the early stages of MI, we conducted further clustering analysis on immune cells from the RAW Data (E-MTAB-7895). As shown in Figure5 -figure Supplement 1A and 1B, seven immune cell types were identified: neutrophils (S100a8 and S100a9), monocytes I (Chil3 and Plac8) and II (Saa3 and Arg1), macrophages (C1qa and Cd68), dendritic cells (H2ab1 and Cd74), B cells (Cd79a and Igkc), and natural killer T cells (Nkg7 and Ms4a4b). The R Package “CellChat” was used to investigate the communication and interaction between these cells and LEC subpopulations. As expected, the communication strength of all significant signaling pathway analyses showed that the highest level of communication probability occured between LEC ca I, NKT and B cells in normal heart tissue, as shown in Figure 5A. During the early phase of MI, that is, between D1 and D3, the highest interaction numbers and strength found were between LEC ca I and ca III, as well as monocytes and macrophages (Figure 5A). Next, correlation analysis was conducted on the strength of the receptor-ligand of LEC and immune cells, except for the classical APP-CD74 pathway (MHCII, expressed by activated APC cells); the interaction was found to be more active in the Galectin9-CD44 and CD45, as well as Collagen-CD44, and ICAM1-ITGβ2 pathways, between the LEC ca I and III, as well as monocytes type I or macrophages (P<0.05) (Figures. 5B and 5C; Figure5-figure Supplement 1C and 1D). This indicates that monocytes and macrophages may be the key immune inflammatory cells that regulate LEC immune presentation during the early phases of MI.

**Figure 5.**
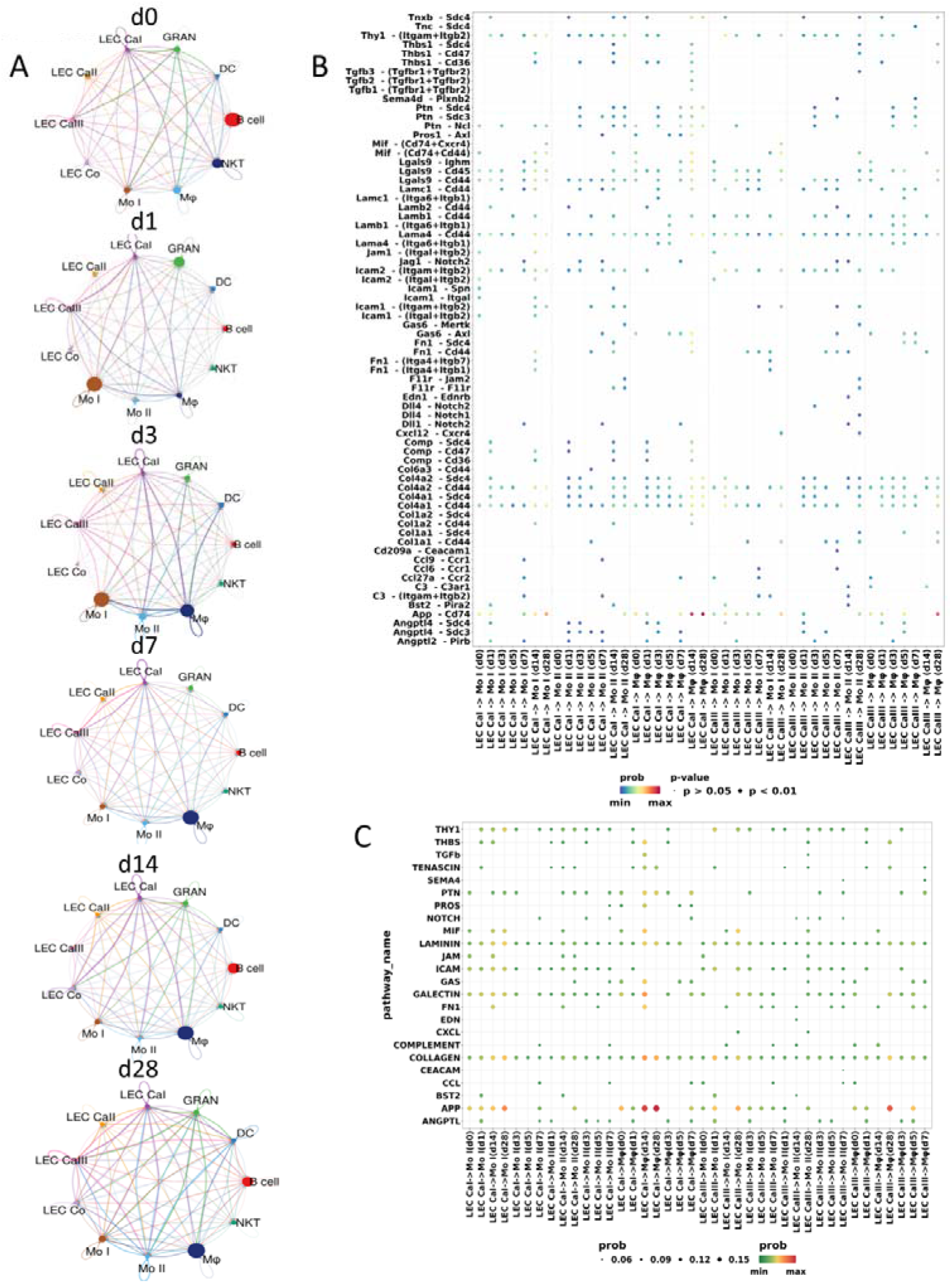
Cell-cell interactions between LECs and immune cells. **(A)** Interaction net count plot of LECs and immune cells. The thickness of the line represents the number of interactions. **(B)** Bubble plots showing the ligand-receptor pairs originating from LEC ca I and ca III that target monocytes and macrophages, the color indicates the probability of interaction and the size indicates the p-value. **(C)** Bubble plots showing the pathways originating from LEC ca I and LEC ca III that target monocytes and macrophages, the color indicates the probability of interaction and the size indicates the p-value.

### 2.6 Aqp1 in LEC is correlated with myocardial edema occurrence and resolution post-MI

In this study, we reveal that fluid retention, one of the basic functions of lymphatic vessels, occurs in LEC ca I and LEC co in the middle-late phase of MI(Figure 3D). Therefore, we explored all molecules that may have important regulatory effects on edema regression. Through localization and temporal expression analysis of KEGG pathway genes in the LEC subpopulation, the expression of Aquaporin 1(Aqp1) in LEC was found to be significantly correlated with temporal changes in edema occurrence and resolution (Figure 6A). Furthermore, the score of the degree of tissue edema in cardiac tissue post-MI, was analyzed and the mouse heart was found to rapidly develop edema on D1 after MI, with the edema reaching its maximum on D3. The edema gradually decreased on D7, and returned to normal on D14 (Figures. 6B and 6C). This was negatively correlated with the expression level of Aqp1 in LEC after myocardial infarction(Figure6-figure supplement 1A). IF co-staining of Aqp1 and LEC in heart tissue also revealed that the expression of Aqp1 was significantly reduced in the early phase of MI D1 and D3, and markedly increased expressed in MI D7 and D14 post-MI (Figure 6D, Figure6-figure supplement 1B). mLECs (mouse LEC cell line) was used and CoCl_2_ intervention (simulated hypoxia stimulation) was found to reduce the expression of Aqp1 at both the transcriptional and expression levels in a dose-dependent manner (Figures. 6E-6J). These results suggest that Aqp1 may be involved in the regulation of lymphatic edema metabolism after MI. Early intervention or increased gene expression may improve cardiac function in MI.

**Figure 6.**
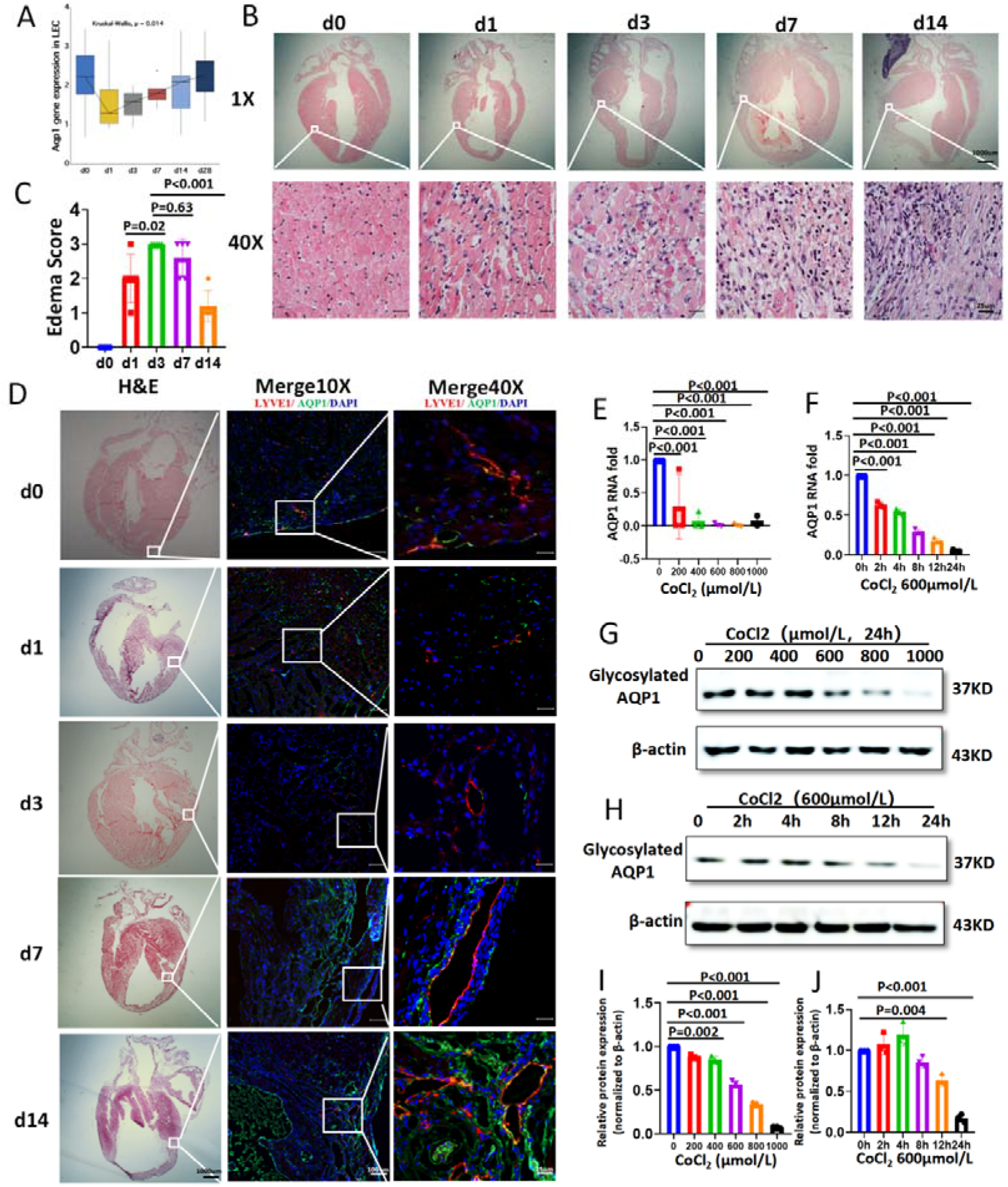
Aqp1 expression on LECs in mouse MI model and on LEC cell line. **(A)** Expression of Aqp1 in LECs at different time points post-MI from the Sc-RNA sequence data. Kruskal□Wallis analysis was performed to compare differences among groups; **(B)**Hematoxylin-eosin(H&E) stained tissue section in sham and MI modle in C57 mouse(n=5); **(C)**Quantification of the edema in infarcted hearts,(n=5); **(D)**H&E stained and immunofluorescence(IF) stained tissue section in sham and MI models in C57 mice(n=5∼7), LYVE1:lymphatic vessel endothelial hyaluronan receptor 1; AQP1: aquaporin1; DAPI: 4’6-diamidino-2-phenylindole; the scale bar is in 10**×**-100μm, 40**×**-25μm. **(E**,**F)**Aqp1 RNA expression in mLEC cell line under CoCl_2_ hypoxic stimulation over different time and different concentrations; (G,H,I,J) Aqp1 protein expression in mLEC cell line under CoCl_2_ hypoxic stimulation over different time pointss and concentrations.

### 2.7 Galectin9 expressed by LEC can affect macrophage migration

To clarify whether the Galectin9(Gal9)-CD44 axis is involved in the regulation of the interaction between LECs and monocyte-macrophages in the early stages of MI, we looked for the temporal expression changes of Gal9 in LECs post-MI. Consistent with the results of the bioinformatics analysis, Gal9 was highly expressed in LECs D3 and D7(Figure 7A), whereas CD44 expression gradually decreased from D1(Figure 7B). Further in vitro studies have found that Gal9 expression was significantly increased in a dose-dependent manner in IFNγ-stimulated cultured mLECs but not hypoxia-stimulated cells (Figures 7C-7E, Figure7-figure supplement 1A-D). Moreover, intervention with recombinant Gal9 significantly induced macrophage migration in a dose-dependent manner (Figures. 7F-7H). In contrast, the inhibition of Gal9 expression significantly inhibited interferon regulated macrophage migration (Figures. 7I-7M). These results further clarify that LECs may participate in the regulation of myocardial tissue immune inflammation through the Gal9-CD44 axis in the early stages of MI.

**Figure 7.**
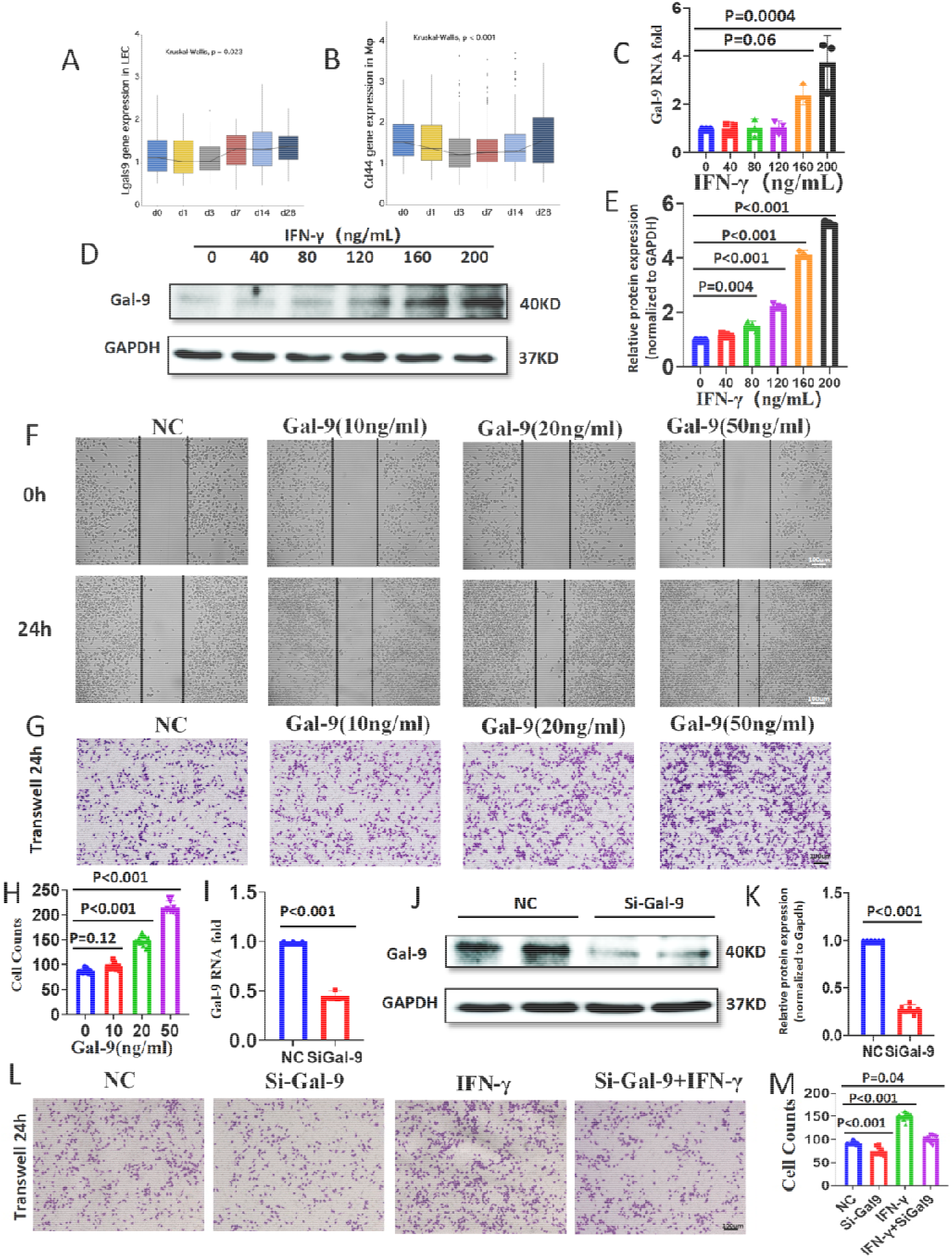
LEC Affect Infiltration of Macrophages Through the Galectin9-CD44 Pathway. **(A and B)** Expression of Lgals9 in LECs and CD44 in macrophages at different time points post-MI from the Sc-RNA sequence data. Kruskal□Wallis analysis was performed to compare differences among groups; **(C)** Gal-9 RNA expression in mLEC cell line under IFN-γ stimulation over different concentration; **(D and E)** Gal-9 RNA expression in mLEC cell line under IFN-γ stimulation in different concentration; **(F)** Mouse monocyte macrophage(J774A.1) wound healing assay under Gal-9 stimulation over different concentration; **(G and H)** J774A.1 macrophage cultured in Transwell system under Gal-9 stimulation over different concentration; **(I, J, and K)** Gal-9 RNA and protein expression after Lgals9 gene.silenced; **(L and M)**J774A.1(in upper cabinet) and LEC (in lower cabinet) cocultured in a transwell system under INF-*γ* stimulation with or without SiGal-9.

## 3 Discussion

Cardiac lymphatics play an important role in myocardial edema and inflammation. This study, for the first time, integrated single-cell sequencing data and spatial transcriptome data from mouse heart tissue at different time points post-MI, and identified four transcriptionally distinct subtypes of LECs and their dynamic transcriptional heterogeneity distribution in different regions of myocardial tissue post-MI. These subgroups of LECs were shown to form different function involved in the inflammation, apoptosis, ferroptosis, and water absorption related regulation of vasopressin during the process of myocardial repair after MI.

The lymphatic system plays an essential role in interstitial fluid balance and immune cell hemostasis in many organs (Petrova & Koh, 2020), including the heart (Brakenhielm & Alitalo, 2019). Stimulating lymphangiogenesis post-MI was found to alleviate myocardial edema and improve heart function (Henri et al., 2016; Houssari et al., 2020; Trincot et al., 2019). Henri et al (Henri et al., 2016) observed lymphangiogenesis and heart function on the 4 or 12-weeks post-MI, and found reduced myocardial edema and fibrosis in VEGFC stimulated mice as compared to the control group. However, lymphangiogenesis in the natural pathological process post-MI, especially in the early-stage post-MI, was not observed in this study. Generally, myocardial edema and inflammation are occurred in MI early stage, what does cardiac lymphatics changed simultaneously caught our interest. So we established mice MI model to observe the physiological and pathological changes in cardiac lymphatics. Interestingly the rate and morphology of lymphatic vessel neogenesis were found to vary in different zone of the infarcted heart, and several large, dilated, continuous Lyve1-labeled lymphatic vessels appeared along the middle myocardium quickly in the border zone in D7 post-MI,which was mainly observed in epicardium and endocardium in the normal heart, this abnormal remolding indicate their new role in this pathological scenario.

Takeda et al (Takeda et al., 2019) have unveiled LEC heterogeneity and their function in human lymphatics, dividing LEC into six subtypes, and found that CD209 on medullary sinuses LECs mediates neutrophil adhesion in the human lymphatic node medulla. Lymphatic vessels LECs, however, have been teeny elaborated. In this study, LECs of lymphatic vessels were clustered into one collecting LEC and three capillary LEC, and, after localized them in space, found that each subtype focuses on different functions in infarcted heart. LEC ca II and III mainly participate in inflammation in the early phase of post-MI in the RZ and IZ, while ca I and co are involved in the regulation of myocardial edema in the middle and late stages after MI in the BZ and IZ. In addition to these traditional roles in immune regulation and interstitial fluid hemostasis, LEC capillaries, especially LEC ca III, also participate in metabolic regulation (glycolysis and gluconeogenesis), which might provide energy for immune cell migrantion or myocardium survival(Guak & Krawczyk, 2020; Zuurbier et al., 2020).

Previous studies have reported that lymphangiogenesis stimulation post-MI can suppress T-cell infiltration and reduce infarct thinning (Houssari et al., 2020), and that activated LECs penetrating the infarcted heart increase the number of tolerogenic dendritic cells and regulatory T cells, thereby inhibiting the expansion of NKT cells and promoting reparative macrophage expansion to facilitate post-MI repair(Wang et al., 2023). This provides a clue for the treatment of lymphatics in infarcted heart.

Herein, we also identified some interesting findings from the Cell-chat analysis between LECs and immune cells. We found that LECs were mainly involved in the chemotactic function of neutrophils, monocytes, and macrophages during the early stages of MI. Macrophage chemotaxis not only participates in the inflammatory response after myocardial infarction, but may also serve as an important immune presenting cell involved in the immune regulation of LEC in myocardial tissue after MI. Second, we found that LEC could regulate monocyte and macrophage infiltration through the Galectin9-CD44 pathway. Galectin9 release from LEC in hypoxia is reduced, which slows macrophage migration, increased Galectin9 in the medium of the macrophage can accelerates its migration. Galectin 9 is one of the β-galactoside binding lectins known for its immunomodulatory role in various microbial infections(Moar & Tandon, 2021), it has been reported that serum Galectin 9 levels are associated with the severity of coronary artery disease, and the ST-segment elevation MI patients have the lowest serum level of Galectin9, followed by the non-ST segment elevation MI group and the stable angina groups(Zhu et al., 2015). Therefore, galectin9 may be a potential molecule involved in regulating the inflammation response post-MI.

Moreover, we screened out a key edema regulated gene, Aqp1, which is highly expressed in LEC subsets in the middle and late phases of MI. Aqp1 is expressed in many tissues, and its primary function is transmembrane water flux(Trillo-Contreras, Toledo-Aral, Echevarría, & Villadiego, 2019). A previous study showed that the knockout of the Aqp1 gene can result in impaired urine concentration(Z. Qiu, Jiang, Li, Wang, & Yang, 2023) and lung edema(Bai et al., 1999) in mouse model. We showed that Aqp1 expression decreased at both the RNA and protein levels in mouse LECs under hypoxic stimulation, and that its expression was negatively correlated with myocardial edema in LECs post-MI. This suggests that Aqp1 may be an essential factor in myocardial edema and needs to be elucidated, along with the underlying mechanism.

In conclusion, LECs play an important role in regulating myocardial edema and inflammation post-MI, and stimulation of lymphangiogenesis or promotion of LEC function may become new targets for future clinical improvement of cardiac function post-MI. One limitation of this study is that the single-cell sequencing data that were analyzed for each time point was derived from a single sample, which resulted in resultant LEC counts being too small. This might have resulted in bias in the analyses. In future studies, multiple samples should be utilized to improve the yield of trace cells like LEC for single cell sequencing. Therefore, further validation tests must be designed and implemented.

## 4 Materials and Methods

### 4.1 Animals and MI model

All animal care and experimental procedures were approved by the Department of Cardiology of the General Hospital of Northern Theater Command and the Cardiovascular Research Institute of the PLA, in accordance with the National Institutes of Health regulations on the use of laboratory animals. Eight-week-old male C57BL/6J mice were obtained from the GemPharmatech Co., Ltd (Nanjing, China). Myocardial infarction Models established by ligation of the left anterior descending artery methodology referred to published study (Liu et al., 2021).

### 4.2 Ultrasound Assessment of Cardiac Function

Ultrasonography was performed using a small animal ultrasound imaging system (Vevo 2100, FUJIFILM VisualSonics, Canada) equipped with a 30 MHz linear ultrasonic transducer. The animals were anesthetized with 2% isoflurane during the ultrasound. The observer who performed the investigation was blinded to the experimental groups. The main cardiac structural and functional measurements were referred to the published study(Liu et al., 2021).

### 4.3 Immunofluorescence and Quantification of Lymphatics

Mice hearts were harvested and postfixed in 4□ 4% paraformaldehyde (PFA) for 4 hours, following dehydration in 4□ 30% sucrose solution overnight, then 4μm frozen slices were sectioned after OCT embedded. Hematoxylin & Eosin staining was performed under standard procedures. For Immunofluorescence(IF), frozen section were blocked with 5% normal goat serum (BioGenex, Fremont, CA) at room temperature for 1hour, and then stained with primary antibodies including, rabbit anti-Lyve1 (1:300, ab14917, Abcam, UK), [Syrian hamster anti-Podoplanin (1:100, 53-5381-82, Thermo, USA), rabbit anti-Prox1(1:300, ab199359, Abcam, UK), both anti-podoplain and anti-Prox1 are additional markers co-stained with Lyve1 to exclude non-lymphatic cells from lymphatic vessel], rabbit anti-CD68 (1:200, CST, 97778S, USA) and mouse anti-Aqp1(1:300, sc-25287, Santa, USA) overnight at 4□, Alexa Fluor 488-or 595-conjugated secondary antibodies including, wheat germ agglutinin (WGA) (1:500, Sigma-Aldrich) in the dark for 1 hours at room temperature on secondary morning. After 5 minutes 4’,6-diamidino-2-phenylindole (DAPI) (Invitrogen USA) staining for cell nuclei, images were captured using a confocal microscope (LSM800; Carl Zeiss Microscopy Ltd, Cambridge, MA).

For quantification of vessel area, vessels with visible co-staining were measured using Image J software. First, we selected an image, turned it into 8-bit, and then applied a suitable threshold adjustment (present co-stained areas wherever possible). ROI manager tools were used next to analyze the automatic signal intensity quantification and AOD(Average Optical Density, %Area) calculated by the software in the area. Finally, the GraphPad software was used to plot the results as a bar graph.

### 4.4 Edema Quantification

Edema quantification from histology images of cardiac paraffin sections were referred to the published study(Trincot et al., 2019). Four high-power (40×) representative images were chosen per animal under the H&E stained section; each image must have a clear border of the section visible. Images were blinded, and five visual fields per sample were evaluated. Subsequently, an edema score was determined for each sample (Score 1=no edema, 2=mild edema, 3=severe edema). Graphs represent the average score value per animal.

### 4.5 Cell culture

The mouse lymphatic endothelial cell line(mLECs) was gifted by Prof. Lin Chen (Army Medical University, China). Mouse monocyte macrophage(J774A.1) was purchased from FuHeng Cell Center (Shanghai, China). All the cell lines were cultured in DMEM with 10% fetal bovine serum(VivaCell Biosciences, China) and 1% antibiotics(penicillin and streptomycin) (Sangon Biothec, China) at 37□.

### 4.6 Cell transfection

Small interfering RNAs(SiRNAs) were purchased from GenePharma (Shanghai, China). The sequences of SiRNA-NC and SiRNA-Lgals9 are listed in **Table S1**. SiRNA were transfected into cells using Transfection Reagent (0000002157, PolyPlus, France) when LEC cell had 40-60% confluence. The efficiency of transfection was detected by RT-qPCR and WB after transfection.

### 4.7 Cell Migration Test

Cell migration was measured by wound healing assay and transwell assay. Wound healing assay procedure were referred to the published study(Martinotti & Ranzato, 2020). For transwell assay, 2.5×10^4^ J774A.1 cells were inoculated into the upper chamber and cultured in serum-free medium. The lower cavity contains 20% fetal bovine serum DMEM medium with or without LEC cocultured. After 24 hours, the migrated cells were immobilized with methanol and stained with 0.1% crystal violet solution (Solarbio, China) at room temperature for 20 minutes. The number of migrating cells was quantified by counting the number of cells in 10 random fields under a 40X magnifying glass.

### 4.8 Western Blotting

mLEC were lysed in RIPA buffer (UJ289235, Thermo Fisher Scientific, USA). Equal amounts of protein were separated by SDS-PAGE (120 V for 1 h) and then transferred onto a polyvinylidene difluoride membrane (R9KA84149, Merck Millipore Ltd. Germany). The membranes were then incubated with the following primary antibodies at 4 °C overnight: mouse anti-Aqp1 (1:1000, sc-25287, Santa, USA), rabbit anti-Gal9 (1:1000, ab69630, Abcam, UK) and rabbit anti-GAPDH (1:1000, CST, USA). After incubation, the membranes were washed and incubated with goat anti-rabbit IgG (1:2000, 150783, Jackson ImmunoResearch, USA) for 1h at room temperature. Finally, the protein bands of the membranes were detected by chemiluminescence with ECL (BIO-RAD, USA). The WB bands were quantitated via the software ImageJ.

### 4.9 Real-Time PCR

Total RNA from mLEC was extracted using the TRIzol reagent (Thermo Fisher Scientific, USA). The isolated RNA was converted into cDNA using the PrimeScript RT with gDNA Eraser kit (Takara, Japan). Quantitative real-time PCR was performed using SYBR1 Premix Ex Taq II (Takara, Japan). The sequences of primers (forward and reverse) are presented in **Table S2**.

### 4.10 Data Collection

Single-cell transcriptomic data was retrieved from ArrayExpress (E-MTAB-7895). Single-cell nucleus transcriptomic data and spatial transcriptomic were retrieved from the Gene Expression Omnibus (GEO; http://www.ncbi.nlm.nih.gov/geo/) database (GSE214611). The animal models used in both E-MTAB-7895 and GSE214611 are permanent MI.

### 4.11 scRNA Analysis

The E-MTAB-7895 data set provided this study with filtered data for 0, 1, 3, 7, 14, and 28 days post-MI. After downloading the matrices, downstream analysis was performed using the Seurat package (version 4.3.0)(Hao et al., 2021) in R (version 4.1.3; R Foundation). Dimension reduction was performed using principal-component analysis (PCA) with the RunPCA function and the optimal number of principal components (PCs) was selected using the ElbowPlot function. The same PCs were also applied to cell clustering with modularity optimization using the kNN graph algorithm as the input. Cell clusters were visualized using the uniform manifold approximation and projection (UMAP) algorithm. The cell types were annotated based on the expression of known marker genes, as previously described. Marker genes for each cell type were obtained using the Seurat function FindAllMarkers, in which only markers with positive log2-transformed fold changes were considered. Differentially expressed genes (DEGs) in each cluster were then identified using the Wilcoxon test.

To characterize the process of LEC cell development and determine lineage differentiation among diverse LEC cells, cell trajectory analysis was conducted using single-cell pseudo-time trajectories constructed with the Monocle2 package (v2.22.0),(X. Qiu et al., 2017) according to the provided documentation (http://cole-trapnell-lab.github.io/monocle-release/). UMI counts were modeled as a negative binomial distribution; ordering genes were identified as having high dispersion across cells (mean_expression≥0.01; dispersion_empirical≥1). The discriminative dimensionality reduction with trees (DDRTree) method was used to reduce data to two dimensions. Gene sets identified from the destination analysis were clustered and visualized using the plot_genes_in_pseudotime function.

The differences in putative cell-cell communication modules between cell types were analyzed for the samples at each stage. The Cellchat (v1.6.1) package includes a comprehensive signaling molecule interaction database that considers the known structural composition of receptor-ligand interactions, such as multimeric receptor-ligand complexes, soluble agonists and antagonists, as well as stimulatory and inhibitory membrane-bound coreceptors(Jin et al., 2021). The inference of cell-cell interactions includes the identification of differentially expressed signaling genes, calculation of ensemble average expression and intercellular communication probability, as well as identification of statistically significant intercellular communications. The single-cell transcriptomic data was divided into seven parts according to the stages and the differences in ligand-receptor interactions and signaling pathways, were explored across these seven stages.

### 4.12 Spatial Localization of LEC Cells

Single nucleus and spatial transcriptome data at 1h, 3 days, and 7days were downloaded (GSE214611). We conducted spatial transcriptome data analysis using the deconvolution algorithm. The deconvolution algorithm refers to the application of feature genes to infer the full matrix information of single-cell transcriptome of cell subclusters. We then compared and anchored the matrix information of the single-cell transcriptome with the information of each SPOT in the spatial transcriptome, predicting cell types based on the similarity between the two sets of information.

### 4.13 Statistic analysis

Data are presented as mean±standard error of mean (SEM). All data were analyzed using SPSS version 22.0 (SPSS, Chicago, IL, USA), and graphs were prepared using GraphPad Prism 8 (GraphPad Software, CA, USA). Differences between two groups were evaluated using an unpaired Student’s t-test. Differences among three or more groups were evaluated using one-way analysis of variance (ANOVA). Statistical significance was set at P<0.05.

## 5 Data availability

Data been analyzed in this study is publicly available from ArrayExpress(E-MTAB-7895) and the Gene Expression Omnibus (GEO; https://www.ncbi.nlm.nih.gov/geo/) database(GSE214611).

## 7 Author Contributions

YLH and CHY contributed to the conception and made final approval of the version; JQH and DLZ performed the study concept, wrote the manuscript, and performed the experiment; HXS, ZQL, DL, XLZ, XJZ, YZ, JL, and JXX helped with the experiment and data analysis. All authors contributed to the manuscript revision, read and approved the submitted version.

## Supporting information

the gene features used to identify in spatial transcriptomics the different LEC subpopulations

