## Supplementary material for "Single-Cell and Spatial Transcriptomic Analyses Reveals the Dynamic Transcript Profiles of Myocardial Lymphangiogenesis post Myocardial Infarction": supplemeantal table1

eLife

Westbrook Centre
Milton Road
Cambridge CB4 1YG, UK

P 01223 855340
W elifesciences.org
T @elife

### Materials Design Analysis Reporting (MDAR) Checklist for Authors

The MDAR framework establishes a minimum set of requirements in transparent reporting mainly applicable to studies in the life sciences.

*eLife* asks authors to **provide detailed information within their article** to facilitate the interpretation and replication of their work. Authors can also upload supporting materials to comply with relevant reporting guidelines for health-related research (see EQUATOR Network), life science research (see the BioSharing Information Resource), or animal research (see the ARRIVE Guidelines and the STRANGE Framework; for details, see *eLife*'s Journal Policies). Where applicable, authors should refer to any relevant reporting standards materials in this form.

For all that apply, please note **where in the article** the information is provided. Please note that we also collect information about data availability and ethics in the submission form.

#### Materials:

| Newly created materials | Indicate where provided: section/figure legend | N/A |
|---|---|---|
| The manuscript includes a dedicated "materials availability statement" providing transparent disclosure about availability of newly created materials including details on how materials can be accessed and describing any restrictions on access. | | No |

| Antibodies | Indicate where provided: section/figure legend | N/A |
|---|---|---|
| For commercial reagents, provide supplier name, catalogue number and RRID, if available. | rabbit anti-Lyve1 (1:300, ab14917, Abcam, UK),<br>rabbit anti-Prox1(1:300, ab199359, Abcam, UK)<br>mouse anti-Aqp1(1:300, sc-25287, Santa, USA)<br>rabbit anti-Gal9 (1:1000, ab69630, Abcam, UK)<br>rabbit anti-GAPDH (1:1000, #5174, CST, USA)<br>rabbit anti-CD68(1:200,97778S,CST,USA)<br>Podoplanin Monoclonal Antibody(1:100,53-5381-82,Thermo,USA) | |

| DNA and RNA sequences | Indicate where provided: section/figure legend | N/A |
|---|---|---|
| Short novel DNA or RNA including primers, probes: Sequences should be included or deposited in a public repository. | RNA oligo sequence of siRNA：<br>siRNA-NC (sense) UUCUCCGAACGUGUCACGUTT<br>siRNA-NC (antisense) ACGUGACACGUUCGGAGAATT<br>siRNA-Lgals9 (sense) GCUCAAACUACCAUCCAUATT<br>siRNA- Lgals9 (antisense) UAUGGAUGGUAGUUUGAGCTT<br>The primer sequences for qRT-PCR：<br>Gapdh(F) TTGACGGAAGGGCACCACCAG | |

| Cell materials | Indicate where provided: section/figure legend | N/A |
|---|---|---|
| Cell lines: Provide species information, strain. Provide accession number in repository OR supplier name, catalog number, clone number, OR RRID. | The mouse lymphatic endothelial cell line (mLECs) was gifted by Prof. Lin Chen (Army Medical University, China). Mouse monocyte macrophage(J774A.1) was purchased from FuHeng Cell Center (Shanghai, China) | |
| Primary cultures: Provide species, strain, sex of origin, genetic modification status. | | No |

| Experimental animals | Indicate where provided: section/figure legend | N/A |
|---|---|---|
| Laboratory animals or Model organisms: Provide species, strain, sex, age, genetic modification status. Provide accession number in repository OR supplier name, catalog number, clone number, OR RRID. | Eight-week-old male C57BL/6J mice were obtained from the GemPharmatech Co., Ltd (Nanjing, China) | |
| Animal observed in or captured from the field: Provide species, sex, and age where possible. | | No |

| Plants and microbes | Indicate where provided: section/figure legend | N/A |
|---|---|---|
| Plants: provide species and strain, ecotype and cultivar where relevant, unique accession number if available, and source (including location for collected wild specimens). | | No |
| Microbes: provide species and strain, unique accession number if available, and source. | | No |

| Human research participants | Indicate where provided: section/figure legend) or state if these demographics were not collected | N/A |
|---|---|---|
| If collected and within the bounds of privacy constraints report on age, sex, gender and ethnicity for all study participants. | | No |

#### Design:

| Study protocol | Indicate where provided: section/figure legend | N/A |
|---|---|---|
| If the study protocol has been pre-registered, provide DOI. For clinical trials, provide the trial registration number OR cite DOI. | | No |

| Laboratory protocol | Indicate where provided: section/figure legend | N/A |
|---|---|---|
| Provide DOI OR other citation details if detailed step-by-step protocols are available. | MI model was referred to PMID: 20966393;Edema quantification was referred to PMI[illegible] | |

| Experimental study design (statistics details) * | | |
|---|---|---|
| **For in vivo studies: State whether and how the following have been done** | **Indicate where provided: section/figure legend. If it could have been done, but was not, write "not done"** | **N/A** |
| Sample size determination | | No |
| Randomisation | | No |
| Blinding | | No |
| Inclusion/exclusion criteria | | No |

| Sample definition and in-laboratory replication | Indicate where provided: section/figure legend | N/A |
|---|---|---|
| State number of times the experiment was replicated in the laboratory. | 3 times | |
| Define whether data describe technical or biological replicates. | | |

| Ethics | Indicate where provided: section/submission form | N/A |
|---|---|---|
| Studies involving human participants: State details of authority granting ethics approval (IRB or equivalent committee(s), provide reference number for approval. | | No |
| Studies involving experimental animals: State details of authority granting ethics approval (IRB or equivalent committee(s), provide reference number for approval. | All animal care and experimental procedures were approved by the Department of Cardiology of the General Hospital of Northern Theater Command and the Cardiovascular Research Institute of the PLA, in accordance with the National Institutes of Health regulations on the use of laboratory animals. | |
| Studies involving specimen and field samples: State if relevant permits obtained, provide details of authority approving study; if none were required, explain why. | | No |

| Dual Use Research of Concern (DURC) | Indicate where provided: section/submission form | N/A |
|---|---|---|
| If study is subject to dual use research of concern regulations, state the authority granting approval and reference number for the regulatory approval. | | No |

#### Analysis:

| Attrition | Indicate where provided: section/figure legend | N/A |
|---|---|---|
| Describe whether exclusion criteria were pre-established. Report if sample or data points were omitted from analysis. If yes, report if this was due to attrition or intentional exclusion and provide justification. | | No |

| Statistics | Indicate where provided: section/figure legend | N/A |
|---|---|---|
| Describe statistical tests used and justify choice of tests. | Data are presented as mean±standard error of mean(SEM). All data were analyzed usi | |

| Data availability | Indicate where provided: section/submission form | N/A |
|---|---|---|
| For newly created and reused datasets, the manuscript includes a data availability statement that provides details for access (or notes restrictions on access). | | No |
| When newly created datasets are publicly available, provide accession number in repository OR DOI and licensing details where available. | | No |
| If reused data is publicly available provide accession number in repository OR DOI, OR URL, OR citation. | Data been analyzed in this study is publicly available from ArrayExpress(E-MTAB-7895) and the Gene Expression Omnibus (GEO; https://www.ncbi.nlm.nih.gov/geo/) database (GSE214611). | |

| Code availability | Indicate where provided: section/figure legend | N/A |
|---|---|---|
| For any computer code/software/mathematical algorithms essential for replicating the main findings of the study, whether newly generated or re-used, the manuscript includes a data availability statement that provides details for access or notes restrictions. | supplementary materials | |
| Where newly generated code is publicly available, provide accession number in repository, OR DOI OR URL and licensing details where available. State any restrictions on code availability or accessibility. | | no |
| If reused code is publicly available provide accession number in repository OR DOI OR URL, OR citation. | | no |

#### Reporting:

The MDAR framework recommends adoption of discipline-specific guidelines, established and endorsed through community initiatives.

| Adherence to community standards | Indicate where provided: section/figure legend | N/A |
|---|---|---|
| State if relevant guidelines (e.g., ICMJE, MIBBI, ARRIVE, STRANGE) have been followed, and whether a checklist (e.g., CONSORT, PRISMA, ARRIVE) is provided with the manuscript. | | |

---

* We provide the following guidance regarding transparent reporting and statistics; we also refer authors to Ten common statistical mistakes to watch out for when writing or reviewing a manuscript.

**Sample-size estimation**

- You should state whether an appropriate sample size was computed when the study was being designed
- You should state the statistical method of sample size computation and any required assumptions
- If no explicit power analysis was used, you should describe how you decided what sample (replicate) size (number) to use

**Replicates**

- You should report how often each experiment was performed
- You should include a definition of biological versus technical replication
- The data obtained should be provided and sufficient information should be provided to indicate the number of independent biological and/or technical replicates
- If you encountered any outliers, you should describe how these were handled
- Criteria for exclusion/inclusion of data should be clearly stated
- High-throughput sequence data should be uploaded before submission, with a private link for reviewers provided (these are available from both GEO and ArrayExpress)

**Statistical reporting**

- Statistical analysis methods should be described and justified
- Raw data should be presented in figures whenever informative to do so (typically when N per group is less than 10)
- For each experiment, you should identify the statistical tests used, exact values of N, definitions of center, methods of multiple test correction, and dispersion and precision measures (e.g., mean, median, SD, SEM, confidence intervals; and, for the major substantive results, a measure of effect size (e.g., Pearson's r, Cohen's d)
- Report exact p-values wherever possible alongside the summary statistics and 95% confidence intervals. These should be reported for all key questions and not only when the p-value is less than 0.05.

**Group allocation**

- Indicate how samples were allocated into experimental groups (in the case of clinical studies, please specify allocation to treatment method); if randomization was used, please also state if restricted randomization was applied
- Indicate if masking was used during group allocation, data collection and/or data analysis
