## Supplemental Table 2 for "Single-Cell and Spatial Transcriptomic Analyses Reveals the Dynamic Transcript Profiles of Myocardial Lymphangiogenesis post Myocardial Infarction"

**Key Resources Table Template and Guidelines**

**Template:**

| **Key Resources Table** | | | | |
|---|---|---|---|---|
| **Reagent type (species) or resource** | **Designation** | **Source or reference** | **Identifiers** | **Additional information** |
| cell line (include species here) | Mouse lymphatic endothelial cell line | gifted by Prof. Lin Chen (Army Medical University, China). | | |
| cell line (include species here) | Mouse monocyte macrophage(J774A.1) | FuHeng Cell Center(Shanghai, China) | FH0329 | |
| transfected construct (include species here) | Transfection Reagent | PolyPlus, France | 0000002157 | |
| antibody | rabbit anti-Lyve1 | Abcam | ab14917 | IF：1:300 |
| antibody | Rabbit anti-CD68 | CST | 97778S | IF:1:200 |
| antibody | Podoplanin Monoclonal Antibody | Thermo | 53-5381-82 | IF:1:100 |
| antibody | rabbit anti-Prox1 | Abcam | ab199359 | IF：1:300 |

| | | | | |
|---|---|---|---|---|
| antibody | mouse anti-Aqp1 | Santa | sc-25287 | IF：1:300<br>WB：1:1000 |
| antibody | rabbit anti-Gal9 | Abcam | ab69630 | WB：1:1000 |
| antibody | rabbit anti-GAPDH | CST | #5174 | WB：1:1000 |
| sequence-based reagent | Aqp1(F) | Sangon Biotech | | ACCTGCTGGCGATTGACTACAC |
| sequence-based reagent | Aqp1(R) | Sangon Biotech | | GTTTGAGAAGTTGCGGGTGAGC |
| sequence-based reagent | Lgals9(F) | Sangon Biotech | | TCTACTCCTGGAATCCCTCCTGTG |
| sequence-based reagent | Lgals9(F) | Sangon Biotech | | AACCTCGTAGCATCTGGCAAGAC |
| peptide, recombinant protein | Galectin-9/lGals9 protein | MedChemExpress | HY-P72637 | |
| peptide, recombinant protein | Interferon-gama | MedChemExpress | HY-P73252 | |
| chemical compound, drug | CoCl2 | Sigma Aldrich | 60818-250g | |
| software, algorith | SPSS | Chicago, IL, USA | | version 22.0 |

| |
|---|
| m |
| other |

4
5
