## Supplementary material for "Single-Cell and Spatial Transcriptomic Analyses Reveals the Dynamic Transcript Profiles of Myocardial Lymphangiogenesis post Myocardial Infarction": Figure supplements

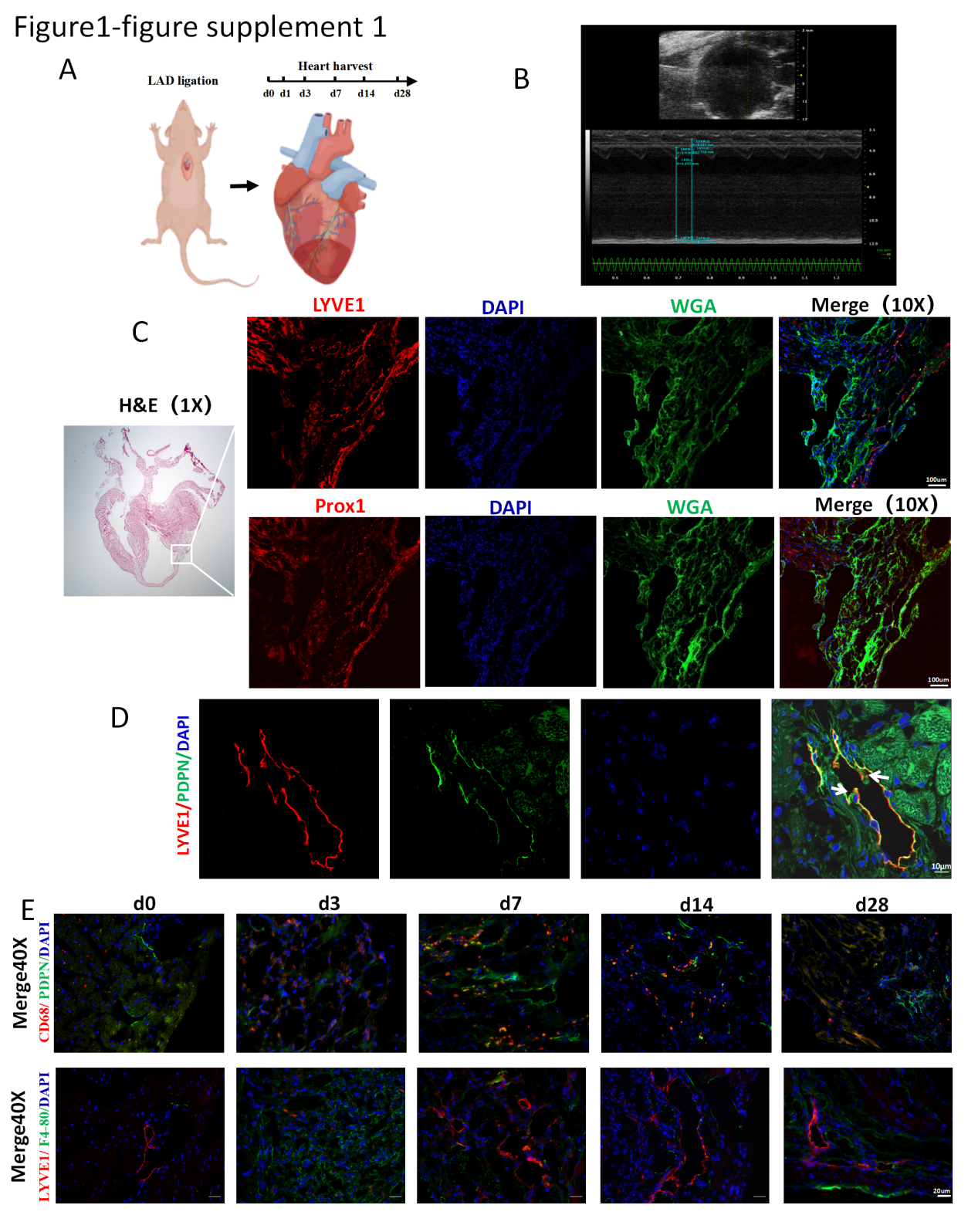

**Figure1-figure supplement 1. (A)** Time points for heart harvest after the establishment of the MI models; **(B)** Transthoracic echocardiography confirmation of the MI mouse model; **(C)** Prox1 and Lyve1 labeled cardiac lymphatics in IF stained frozen tissue section; **(D)** LYVE1&PDPN labeled collecting lymphatics with the lymphatic valves highlighted with white arrows; **(E)** CD68 & Pdpn and LYVE1 & F4-80 co-stained the BZ in different time point. LYVE1: lymphatic vessel endothelial hyaluronan receptor 1; WGA: wheat germ agglutinin; PDPN: Podoplanin; DAPI: 4’6-diamidino-2- phenylindole; scale bar in 1×-1000 μm, 10×-100 μm, 40×-25 μm.

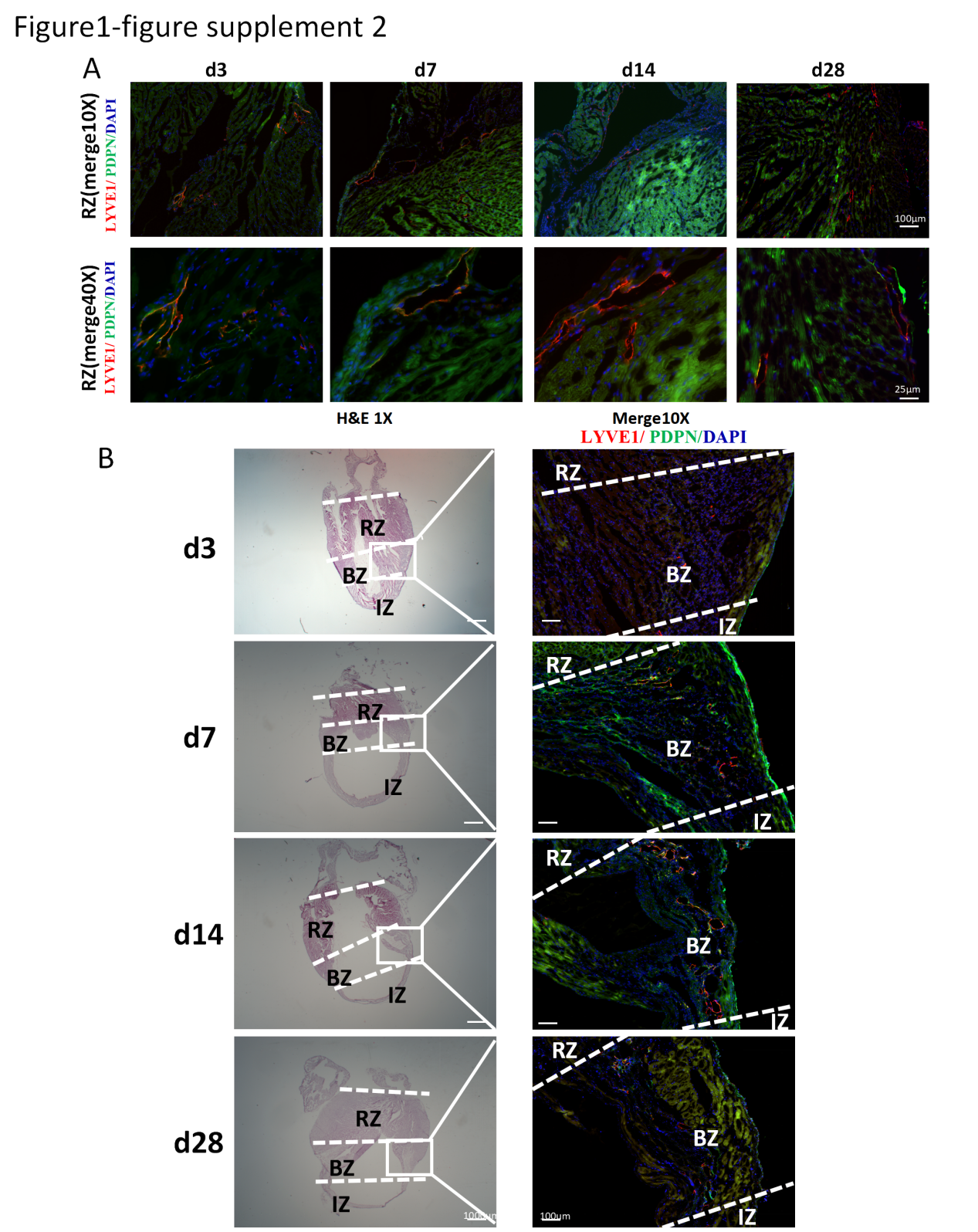

**Figure1-figure supplement 2. (A)** LYVE1 & PDPN co-stained cardiac lymphatics in remote zone of MI models in mouse heart; **(B)** Different Zones are labeled in Hematoxylin-eosin (H&E) staining and immunofluorescence (IF) images in different time points after MI. LYVE1: lymphatic vessel endothelial hyaluronan receptor 1; PDPN: Podoplanin; DAPI: 4’6-diamidino-2- phenylindole; IZ: Infarct Zone; BZ: Border Zone; RZ: Remote Zone; scale bar in 1×-1000 μm, 10×-100 μm, 40×-25 μm.

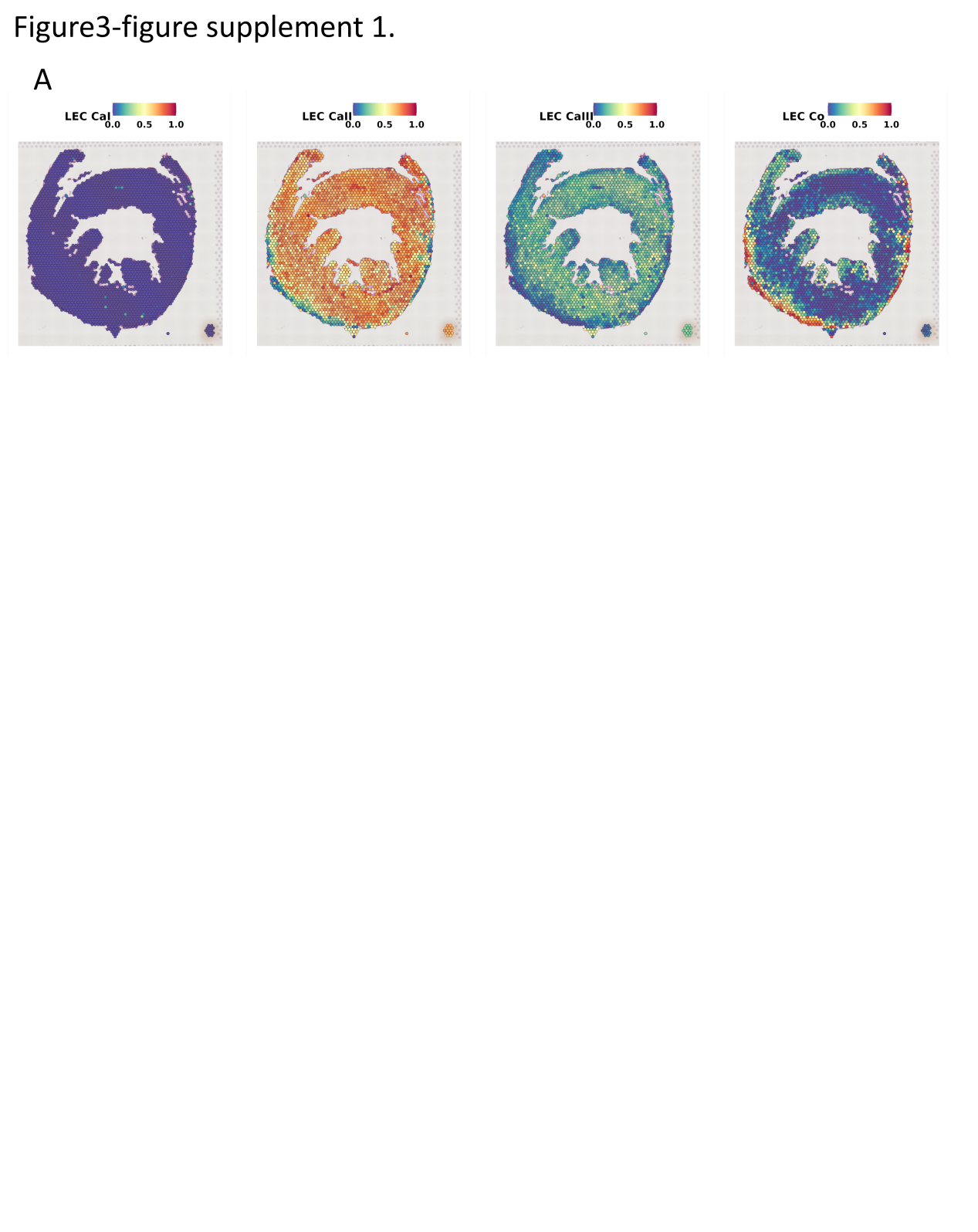

**Figure3-figure supplement 1.** (A) The proportion of LEC cell types in each spot was determined by spatial transcriptome in 1 h post MI (Because we could not download spatial transcriptome data for day d0 in the public database (GSE214611) or from the authors, we have used data of 1 h after IR as a reference for approximating the physiological state).

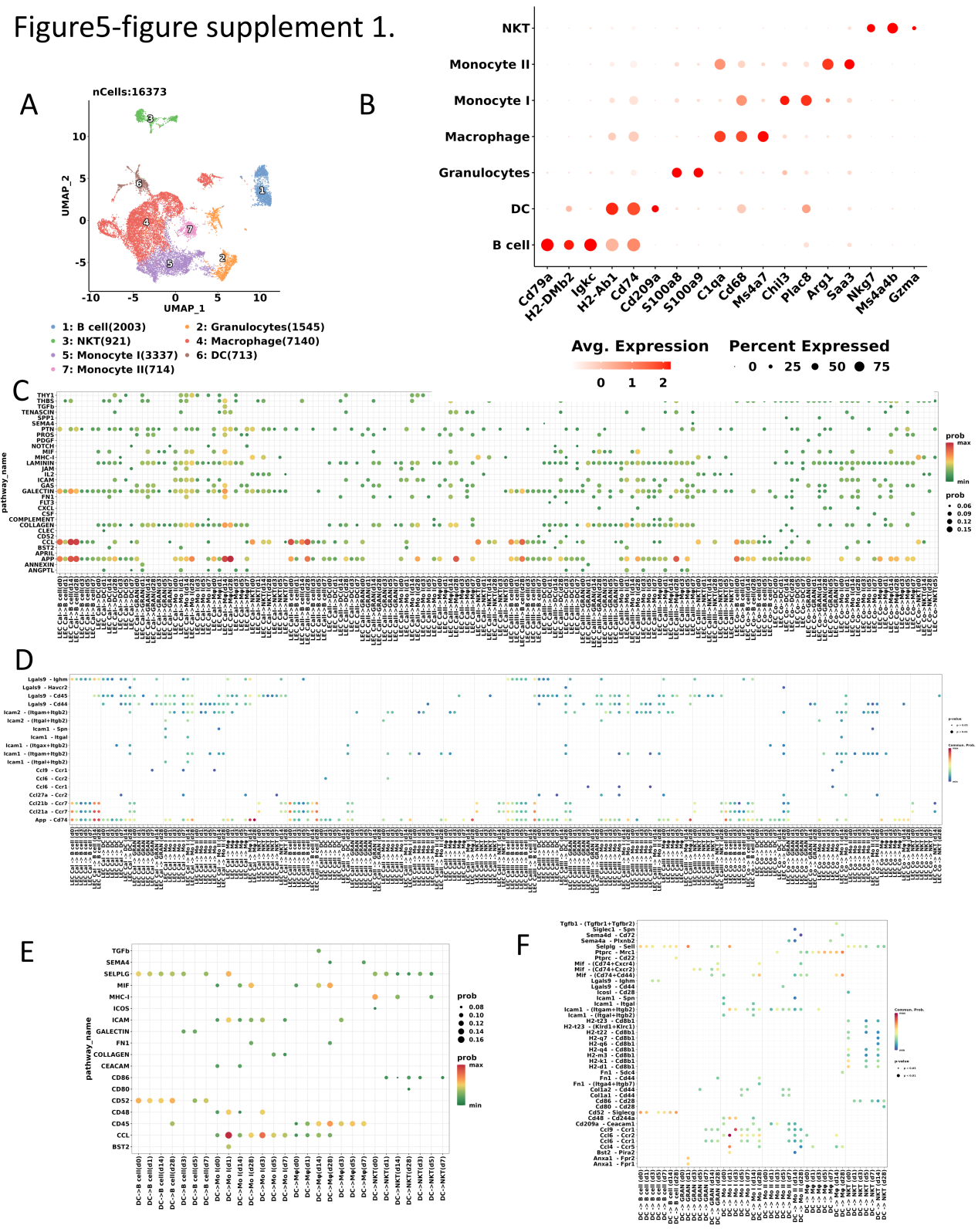

**Figure5-figure supplement 1.** **(A)** A Uniform Manifold Approximation and Projection (UMAP) plot of 16373 cells showing seven identified cell types in immune cells, cell types have been coded with different colors. **(B)** A dot plot showing the average expression of the selected cell marker genes. Dot size and color indicate the percentage of marker gene expression and average expression level (Avg. Expression). **(C)** Bubble plots showing the pathway originating from LECs targeting immune cells; the color indicates the probability of interaction. **(D)** Bubble plots showing the ligand-receptor pairs originating from dendritic cells (DCs) targeting immune cells; the color indicates the probability of interaction and the size indicates the p-value. **(E)** Bubble plots showing the pathways originating from dendritic cells (DCs) targeting other immune cells; color indicates the probability of interaction. **(F)** Bubble plots showing the ligand-receptor pairs originating from dendritic cells (DCs) targeting other immune cells; the color indicates the probability of interaction and the size indicates the p-value.

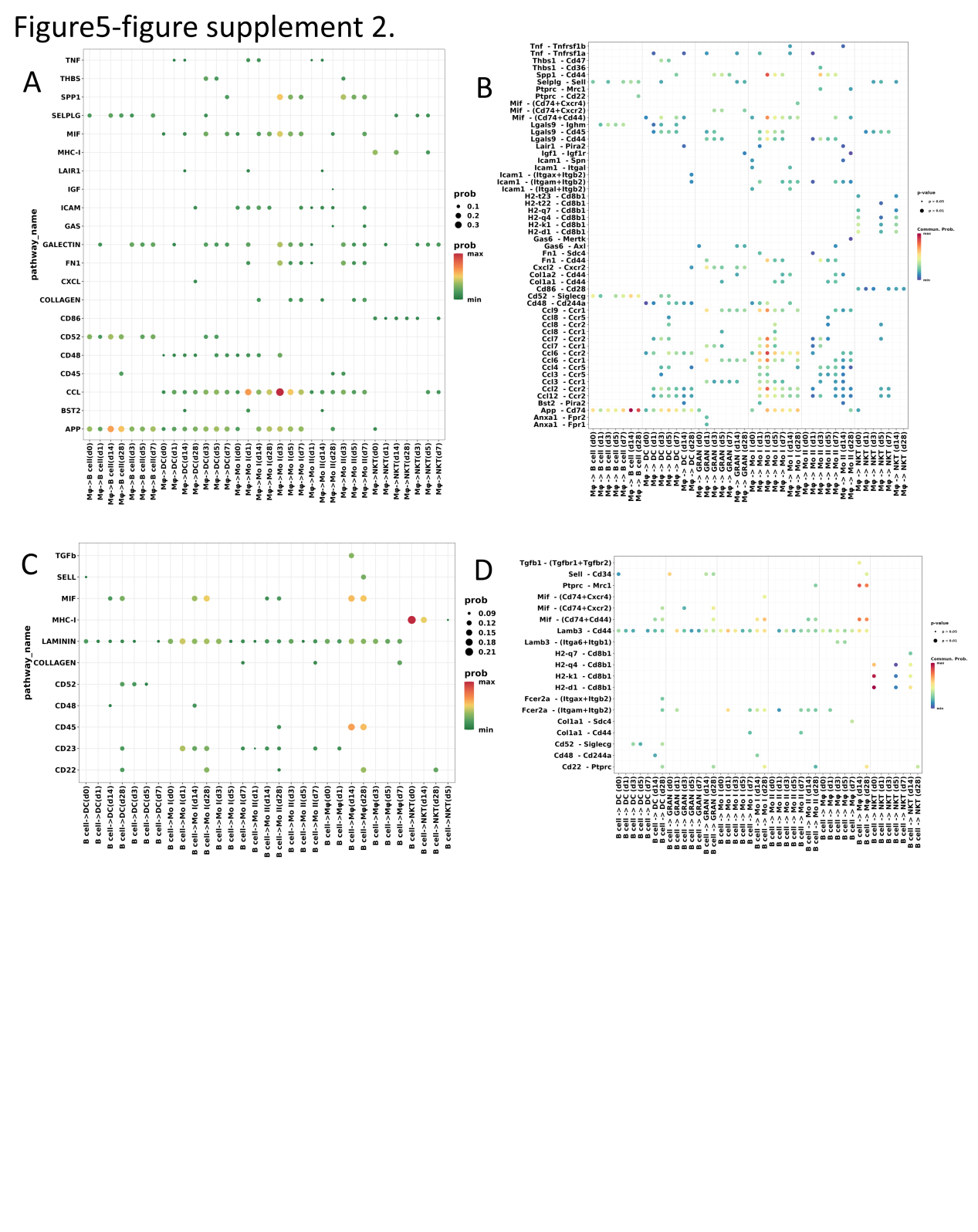

**Figure5-figure supplement 2. (A)** Bubble plots showing the pathways originating from the macrophages (Mφ) targeting other immune cells; the color indicates the probability of interaction. **(B)** Bubble plots showing the ligand-receptor pairs originating from Mφ targeting other immune cells; the color indicates the probability of interaction and the size indicates the p-value. **(C)** Bubble plots showing the pathway originating from B cell targeting other immune cells; the color indicates the probability of interaction. **(D)** Bubble plots showing the ligand-receptor pairs originating from B cells targeting other immune cells; the color indicates the probability of interaction and the size indicates the p-value.

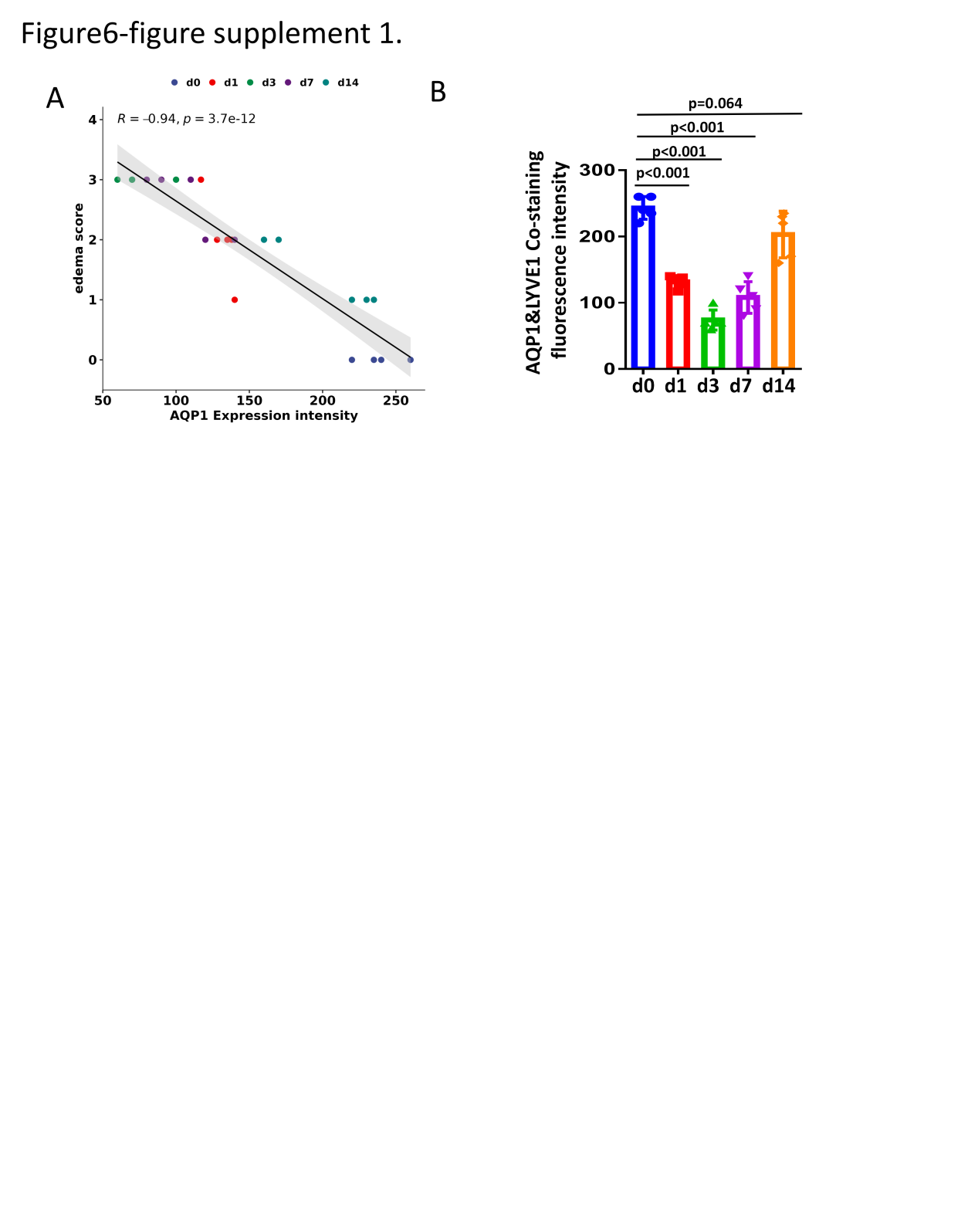

**Figure6-figure supplement 1.** **(A)** Correlation between Aqp1 expression intensity and edema score.; (B)Semi-quantification of AQP1 protein expression in LECs by measure AQP1&LYVE1 co-staining fluorescence intensity.

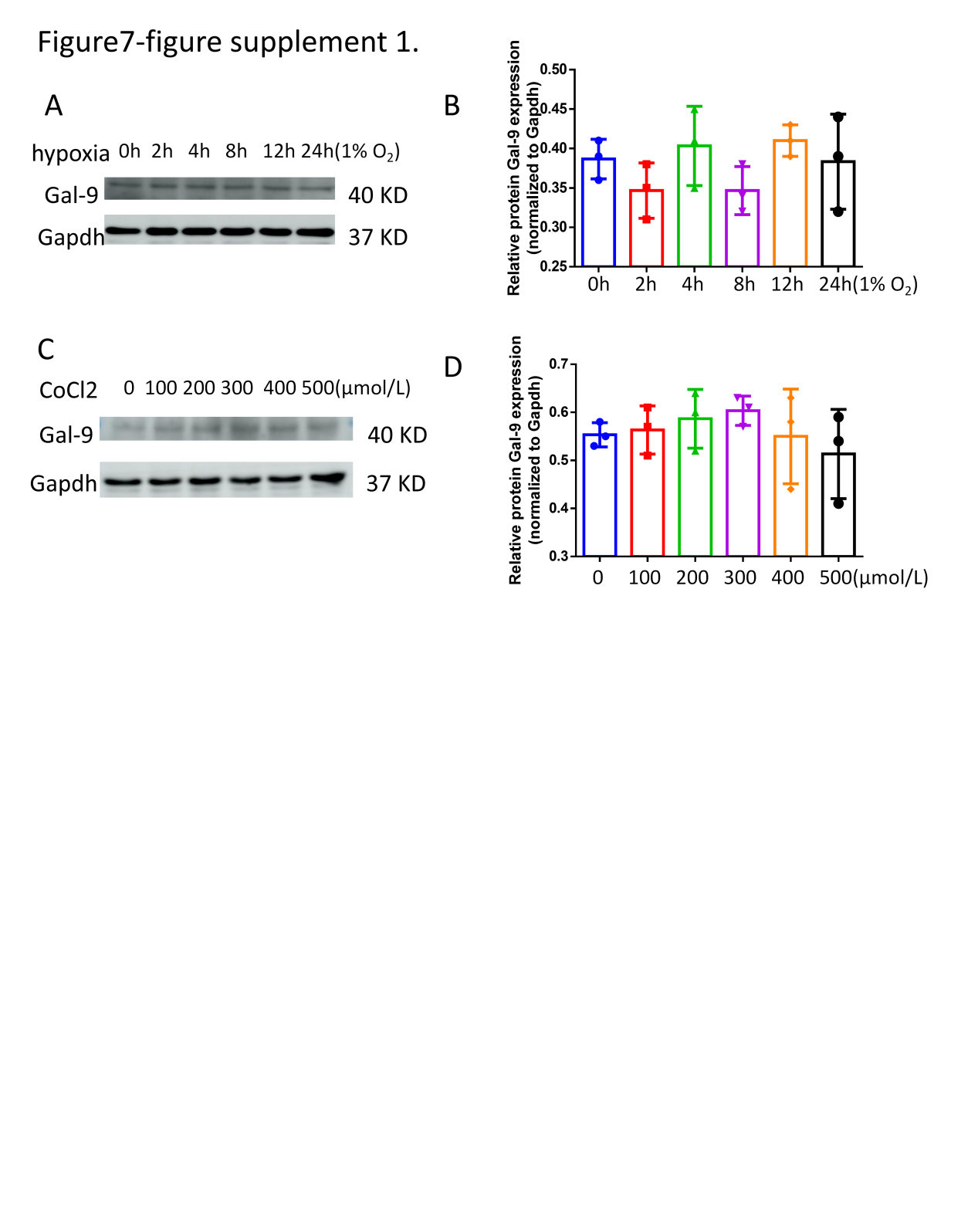

**Figure7-figure supplement 1.** **(A and B)** Expression of Gal-9 protein of LEC under 1% oxygen concentration stimulation over different times; **(C and D)** Expression of Gal-9 protein of LEC under CoCl_2_ intervention over different concentration. By one-way ANOVA test, the protein expression of Gal-9 in LECs stimulated by hypoxia over different times or different concentrations of CoCl_2_ were all not statistically different from the control group(P>0.1).
